# DNA metabarcoding using new *rbcL* and *ITS2* metabarcodes collectively enhance detection efficiency of medicinal plants in single and polyherbal formulations

**DOI:** 10.1101/2023.01.11.523696

**Authors:** Tasnim Travadi, Abhi P. Shah, Ramesh Pandit, Sonal Sharma, Chaitanya Joshi, Madhvi Joshi

**Affiliations:** Gujarat Biotechnology Research Centre (GBRC), Department of Science and Technology, Government of Gujarat, Gandhinagar- 382010, Gujarat, India

**Keywords:** Adulteration, DNA metabarcoding, Herbal medicines, Next Generation sequencing, Pharmacovigilance

## Abstract

With the widespread adoption of barcoding and next-generation sequencing, metabarcoding is emerging as a potential tool for detecting labelled and unlabelled plant species in herbal products. In this study, newly designed *rbcL* and *ITS2* metabarcode primers were validated for metabarcoding using in-house mock controls of medicinal plant gDNA pools and biomass pools. The applicability of the multi-barcode sequencing approach was evaluated on 17 single drugs and 15 polyherbal formulations procured from the Indian market. The *rbcL* metabarcode demonstrated detection efficiencies of 86.7% and 71.7% in gDNA plant pools and biomass mock controls, respectively, while the *ITS2* metabarcode demonstrated 82.2% and 69.4%. In the gDNA plant pool and biomass pool mock controls, the cumulative detection efficiency increased by 100% and 90%, respectively. A total of 79% cumulative detection efficiency of both metabarcodes was observed in single drugs, while 76.3% was observed in polyherbal formulations. An average fidelity of 83.6% was observed for targeted plant species present within mock controls as well as in herbal formulations. Our results demonstrated the applicability of multilocus strategies in metabarcoding as a potential tool for detecting labelled and unlabelled plant species in herbal formulations.

## 1. Introduction

The herbal commodity market sector is thriving, owing to the widespread belief that traditional medicine is natural and thus safer, thereby promoting good health and sustainable life policies. But as the market expands, the shortage of genuine resources and lack of taxonomic knowledge challenge the authenticity of herbal drugs and increase the incidence of economically motivated or unintentional adulterations or substitutions^1^. To retain the trust and safety of consumers and their health, strict pharmacovigilance is a necessity. However, the regulatory guidelines for medicinal plants often blur the line between foods and therapeutics and vary from nation to nation. To close this research gap, regulatory bodies must implement more reliable, universal, and robust detection methods^2^.

Nowadays, various pharmacopoeia is advocating DNA based methods such as DNA barcoding and species-specific PCR assay to authenticate herbal raw material as DNA is more stable, unaffected by the external factors, and present in almost all plant tissues invariably^3^. In addition, the DNA based results are independent of seasonal variations, age of the plant, which in case of chemical marker based methods vary significantly^4^. Therefore, the results of DNA based methods are free from subjectivity, accurate and provide a universally accepted platform for the authentication of botanicals in a wide range of food and herbal products^5^. The advent of DNA barcoding is the first step in this direction, as barcoding gives resolution up to species level^6^. There are 17 potential barcode regions (*matK, rbcL, ITS, ITS2, psbA-trnH, atpF-atpH, ycf5, psbKI, psbM-trnD, coxI, nad1, trnL-F, rpoB, rpoC1, atpF-atpH*, and *rps16*) for plants, having different degrees of universality, specificity, and taxa resolution power that extensively used in the authentication and identification of medicinal plants^4,7^. However, DNA barcoding^8^ and species-specific assays^9–11^ cannot resolve presence of multiple plant species in a single sample^12^. which can be overcome by DNA metabarcoding.

DNA metabarcoding combined the strengths of next generation sequencing and barcoding for detecting multiple taxa in samples^13^. It has proven challenging to use a single plant barcode for species-level identification due to the great diversity, relatively slow molecular evolution, and frequent cross-pollinations and hybridization in the plant kingdom; henceforth, different barcodes show different degrees of taxon specificity^14^. In order to precisely identify the plant species in the sample, multi-barcode approaches have become more prevalent. Xin et al.^15^ and Frigerio et al.^16^ employed multi-barcode approach of *ITS2* and *trnL* for various Chinese medicine and herbal teas. However, these studies also highlighted the limitations of DNA metabarcoding applications for authentication of herbal products due to variability in degrees of universality and resolution power of barcodes for specific taxa, a lack of a curated database, and robust bioinformatics pipeline. To overcome these constraints, there is a need for screening of new barcodes and new variable regions within the same barcode for authentication of the herbal products.

Therefore, the aims of the present study were: 1) to develop a new *rbcL* and *ITS* metabarcode primers for the detection of medicinal plant species 2) to validate the primers efficiency using mock controls and then apply the same for the market formulations (17 different single drugs and 15 polyherbal market formulations). 3) to see whether a multi-barcoding approach could be used to detect targeted plant species in herbal formulations?

## 2. Materials and Methods

### 2.1. Collection of reference plant material and herbal products

Reference plant materials were collected with the aid of a taxonomist from the Maharaja Sayajirao University (MSU), Vadodara (Gujarat, India) and the Directorate of Medicinal and Aromatic Plants Research (DMAPR), Anand (Gujarat, India). Reference plant materials were authenticated by Sanger sequencing of *rbcL* gene as described earlier^17^ and sequences were submitted to the NCBI database (accession number MW628906 to MW628936). Voucher specimens were developed and deposited in our institutional herbarium.

In total, 32 herbal products were collected by blind sampling from the local market and e-commerce, with 17 single drugs and 16 being polyherbal formulations. Single drugs include four Tulsi powder, five Gokhru powder, three Shatavari powder, two Vasa products, and one each of Bhringraj, Ashawgandha, and Arjuna powder. Polyherbal formulations include, three market samples of Trikatu (has three plant species), three samples of Sitopladi [comprises five constituents, only three of which are plant species; the other two are sugar and Vanshlochan (the female bamboo exudate), hence these two constituents were not considered while analyzing the data expecting absence of DNA for these two], four samples of Rasayana (has three plant species), four samples of Hingwashtak (has seven plant species), and one sample of Talisadi [comprises eight constituents, only six of which are plant species; the other two are sugar and Vanshlochan (the female bamboo exudate), hence these two constituents were not considered while analyzing the data expecting absence of DNA for these two].

### 2.2. Primer designing

To design the metabarcodes for *ITS2* gene, ITS2 sequences of magnoliophyta from the BOLD database were downloaded and curated, particularly for the length. For the *rbcL* gene, we used 1,776 sequences of our *in-house* sequencing project that were submitted to the BOLD database. In order to design universal barcodes, *rbcL* gene sequences were filtered by length between 450-600 bp. So finally, 1,465 and 1,701sequences for *ITS2* and *rbcL*, respectively were obtained. These sequences were preceded for multiple sequence alignment separately (*ITS2* and *rbcL*) using BioEdit 7.2. HYDEN (HighlY DEgeNerate primers) software^18^ to design degenerate primers, where, maximum 3 degeneracy per primer were allowed. The designed primers were checked for amplicon length using NCBI primer BLAST^19^. *RbcL* reverse primer sequence was designed in this study, while forward primer sequence was obtained from Maloukh et al.^20^. To synthesized fusion primers, forward primers of *rbcL* and *ITS2* were tagged with the Ion torrent adapter and a 10 bp multiplex identifier barcode, while reverse primers were tagged with the P1 adapter. Nucleotide sequence of the designed primers and their amplicon length are not shown here.

### 2.3. PCR Optimization and library preparation

Library preparation process became a single step process with barcoded fusion primers. The PCR optimization with each barcoded fusion primer was done with 45 different plant DNA listed in Supplementary Information Table S1. Thermal cycler conditions, especially primer annealing temperature, were optimized for *rbcL* and *ITS2* primer pairs with the following conditions. PCR mixture containing 10 µL Emerald Master mix (2X) (TaKaRa), 2 µL total genomic DNA (10-15 ng/µL), 1 µL of forward (5 pmol), 1 µL of reverse primer (5 pmol), 1 µL BSA (2 mg/mL) and 5 µL PCR grade water with the following thermal cycling conditions. Initial denaturation 95°C for 5 minutes followed by 30 cycles of 95°C for 1 minute, for primer annealing a temperature gradient of 50 to 60 °C with interval of 2 °C for 30 seconds and 72°C for 1 minute, and final extension 72°C for 5 minutes.

### 2.4. Preparation of different mock controls

Three different types of controls were prepared as follows: Control 1) genomic DNA from plant leaves from different genus has been first isolated and pooled into different three groups as mentioned below, Control 2) simulated plant biomass controls (blended formulations) in which respective plant part having medicinal value has been mixed together and subjected to DNA isolation and Control 3) genomic DNA (Isolated from plant leaves) pool from different species of the two genus (Fig. 1). As mentioned above, for the first type of control, three different group was prepared having the plant species of different genus. Group one comprised DNA of five species in equal proportion (5P) and further in group 2 and 3 DNA was added from ten (10P) and fifteen different plant species (15P) (Fig. 1, Fig. 2a). High quality DNA of all the species have been isolated individually and pooled together in equal proportion to make these groups. These controls were considered as positive controls, hypothesizing that all the plant species get amplified and resolution power of the primers designed for the study should be free from any biases. Diversity of the group has been increased by adding species from diverse genera covering the wide range in order to test the resolution capacity of the primers for the maximum number of species. For the second type of control, the same three groups of plants that were used in first controls (labelled as 5S, 10S, 15S), but here simulated blended plant parts containing bioactive therapeutics were mixed in equal proportion (biomass admixture controls). To comprehend biases in the DNA extraction process, PCR and the impact of secondary metabolites on PCR amplification, these controls can be used. For the third type of control, two groups were prepared. One group comprises six plant species of the two different genus including *Asparagus* and *Terminalia* (Fig. 2b). Second group comprises seven plant species of the two genus including *Piper* and *Phyllanthus* (Fig. 2b). Similar to the first control, here also high quality DNA that was individually isolated from each species and then pooled in an equal amount. These controls were utilised to obtain insight into the resolving strength of our newly designed *rbcL* and *ITS2* metabarcodes at genus and species level.

**Figure 1.**
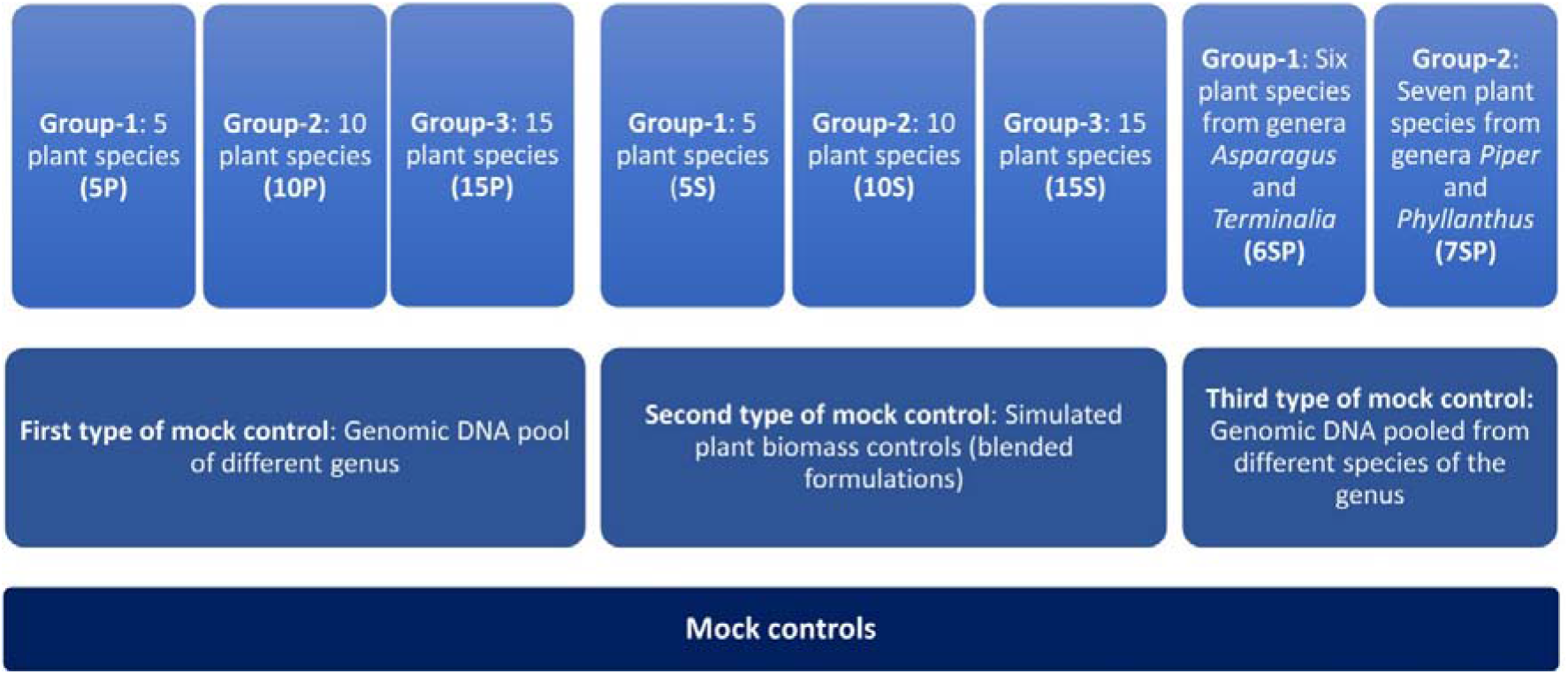
Schematic representation of different types of mock controls prepared in this study. First and second type of mock controls have 3 different groups comprising 5, 10 and 15 plant species. Third type of mock control has 2 different groups, one with a gDNA pool of six plant species from genera *Asparagus* and *Terminalia* and another with a gDNA pool of seven plant species from genera *Piper* and *Phyllanthus*.

**Figure 2.**
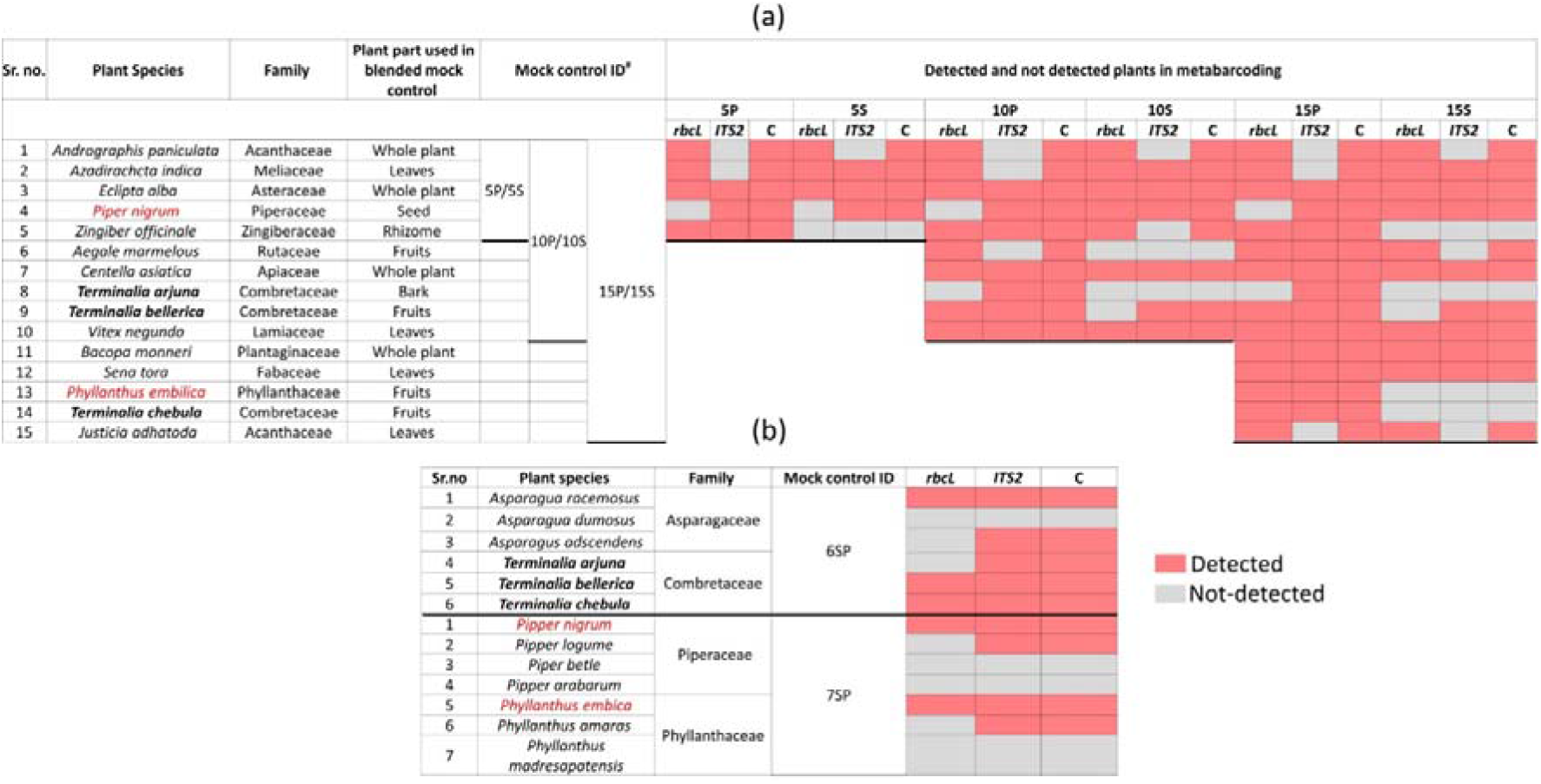
The distribution of predefined herbal species detected in each mock control with *rbcL, ITS2* and combined metabarcoding approach. (a) Detected and undetected plant species with *rbcL, ITS2* and combined approach in first type of mock control i.e. genomic DNA pool of different genus and second type of mock control i.e., simulated plant biomass control (blended formulations) comprising three groups having five plant species (5P/5S), ten plant species (10P/10S) and 15 plant species (15P/15S). (b) Detected and undetected plant species with *rbcL, ITS2* and combined approach in a third type of mock controls (i.e., genomic DNA pooled from different species of the genus) comprising two different groups, one having six plant species (6SP) from genera *Asparagus* and *Terminalia* and another having seven plant species (7SP) from genera *Piper* and *Phyllanthus*. Detected plant species are represented with pink color while undetected plant species are represented in grey color. Plant species, their family, plant parts used in the second type of mock controls (simulated plant biomass controls or blended formulations), mock control IDs also indicated in figure. P: gDNA plant pool controls, S: simulated blended plant pool (biomass control), C: combined approach.

### 2.5. Metabarcoding

DNA from plant materials, blended formulations and herbal products was extracted in duplicate using DNeasy Plant Mini Kit (QIAGEN, Germany) by following the manufacturer’s instructions. Library was prepared from each DNA sample using *rbcL* and *ITS2* fusion primers with optimized PCR conditions. The libraries were purified using AMPure XP beads (Beckman Coulter, CA, USA) and quality of some of the libraries was checked using Agilent high sensitivity DNA kit on Agilent 2100 Bioanalyzer. For each sample, libraries from two replicates were pooled. Further, all libraries were diluted to 100 pM and pooled in equimolar concentration. Emulsion PCR was carried out using Ion 520™ & Ion 530™ Kit-OT2 with 400 bp chemistry (Thermo Fisher Scientific, MA, USA). Sequencing was carried out on the Ion S5 system using a 520/530 chip (Thermo Fisher Scientific, MA, USA).

### 2.6. Metabarcoding data analysis

For establishing the metabarcoding data analysis pipeline, optimization was done with three parameters as follows. 1) filtering criteria includes, discarding reads with length <280 and >350 bp, <300 and >350 bp, <320 and >350 bp for *rbcL* and for *ITS2* discarding reads <280 and >300 bp, 2) OTU clustering with 97, 98 and 99% similarity in reads, 3) discarding OTU clusters having <5 or <10 reads. Obtained reads were filtered based on the quality score (Q >=25 for *rbcL* and Q >=20 for *ITS*) and read length using PRINSEQ^21^. Clustering of filtered reads were performed using CD-HIT-EST^22^. After that, taxonomic assignment of OTU clusters having ≥5 or ≥10 reads were done using BLASTn^19^ (NCBI) with minimum E value 10E^-5^. For each sequence, ten hits were retrieved, and each hit was inspected and evaluated manually for the assigning plant genus and species. To analyse read abundance of each plant species, number of reads were normalised by considering total numbers of reads obtained after discarding clusters having a <5 reads or <10 reads as 100%. Detection efficiency (%) for both metabarcodes was calculated using the following formula. (Total number of detected targeted or labelled plant species /total number of plant species present in herbal formulation) × 100. Fidelity of detection (absolute) can be defined as the total number of samples or herbal formulations in which targeted plant species were detected per total number of samples or herbal formulations in which targeted plant species were present^23^. According to that, relative fidelity of detection (%) was calculated based on the following formula. (Total number of samples or herbal formulations in which targeted plant species detected/total number of samples or herbal formulations in which targeted plant species are present) × 100. Fidelity of detection (absolute or relative) was calculated only where plant species that present in more than one group of the same type of control (n>1) and for market samples sample size is more than one (n>1).

## 3. Results and Discussion

Coghlan et al.^13^ first introduced a metabarcoding approach for the detection of plant and animal raw materials used in 15 traditional Chinese medicines using a p-loop region of the plastid *trnL* gene and 16S mtDNA marker respectively. Later on, several studies reported application of metabarcoding for authentication and detection of plant materials in herbal medicines with single and multi-barcode approach. For instance, Yao et al.^24^ and Cheng et al.^25^ employed multi-barcode approach of *ITS2* and *trnL* in metabarcoding for detection of plant species in various traditional Chinese medicine (TCM). Urumarudappa et al.^26^ used *ITS2* and *rbcL* barcode in metabarcoding for detection of plant species in herbal medicines of Thailand. Using *ITS1* and *ITS2* barcodes in metabarcoding, Raclariu et al.^27–29^ reported presence of unlabelled species by 89, 68, and 15% in single drugs of *Echinacea* species, *Hypericum perforatum*, and *Veronica officinalis*, respectively sold in the European market.

In 2014 and 2015, the total commercial market for herbal material in India was estimated to be more than 512,000 tonnes, with a market value of $1 billion USD^30^. India has over 8,000 authorised medicinal product manufacturing units, and the market growth for herbal products is outstripping supply capacity for some plant species^30^. However, till date detection of raw plant materials of Indian marketed herbal medicine using metabarcoding approach is not well established. Ichim^31^ demonstrated 31% adulteration in 752 Indian marketed herbal products with DNA barcoding and species specific marker based approach but not via metabarcoding. Earlier we have reported presence of unspecified plant species in four polyherbal formulations of the Indian market using *rbcL* minibarcode via metabarcoding approach^17^. Here, we have used a multi-barcode strategy to identify raw plant components in single drugs and polyherbal formulations of the Indian market using newly designed *rbcL* and *ITS2* metabarcodes.

### 3.1. PCR assays using newly designed rbcL and ITS2 primers and fusion primers

Minimal criteria, such as routine amplifiabilities and minimum intraspecific but maximum interspecific divergence at the taxon level, must be followed in the search for the appropriate barcode region. Hence, degenerated *rbcL* and *ITS2* metabarcode primers were designed to get high amplification efficiency, universality, and resolution power. In total 45 medicinal plants from diverse families, genus and species were taken to confirm and optimize the newly designed *rbcL* and *ITS2* primer sets for PCR amplification experimentally (Table S1). The annealing temperature was optimised, and the results showed that the *rbcL* and *ITS2* primer sets performed optimum at a temperature of 56 °C (data not shown). Among newly designed *rbcL* and *ITS2* metabarcodes, *rbcL* has been found to be very robust, universal and gives a 100% amplification efficiency within selected 45 plants, but *ITS2* metabarcode gives 88.9% amplification efficiency and was not able to give amplification in 5 plant species include *Ailanthus excelsa, Andrographis paniculata, Adhatoda vasica, Ocimum tenuiflorum*, and *Ocimum canum* (Table S1). However, due to greater species level discrimination power of *ITS2* in medicinal plants^24,32^, here *ITS2* metabarcode and *rbcL* metabarcode were taken. The amplification effectiveness of “fusion primers” (tagged with Ion torrent adapter and barcodes) of *rbcL* and *ITS2* metabarcodes remained unchanged. However, the appearance of non-specific amplification in some barcodes suggest that 56 °C annealing temperature is not optimal for the fusion primers and therefore, further optimization of annealing temperature revealed that non-specific amplification was overcome by increasing the annealing temperature to 60 °C (data not shown).

### 3.2. Establishing data analysis pipeline using mock controls

For establishing the metabarcoding data analysis pipeline, the first type of mock control was used i.e., gDNA pooled controls of different genus. The first parameter is filtering criteria, for *ITS2* the best filtering criteria was to remove reads with length < 300 bp and for *rbcL* descarding reads with length <300 and >350 bp (data not shown here). The second parameter is the OTU percentage similarity of clustering the reads, in which in case of *rbcL* metabarcode, a greater number of plant species was detected when the reads were clustered at 99% identity. In the case of *ITS2* metabarcode, clustering the reads at 97% and 98% similarity were equally capable for resolving the plant species (Supplementary Information Fig. S1). Therefore, for *ITS2* metabarcode, henceforth, for OTU clustering i.e., 98% with a greater percentage was selected. The third parameter is the discarding OTU clusters having <5 or <10 reads, in which we observed that discarding OTU clusters having <5 reads was able to detect the greater number of plant species in the case of both metabarcodes (Supplementary Information Fig. S2). Based on these findings, while selected read lengths were between 300 to 350 bp, 99% OTU clustering, and discarding of OTU clusters having <5 reads for the *rbcL* metabarcode and read lengths of >300 bp, 98% OTU clustering, and discarding of OTU clusters comprising <5 reads for the *ITS2* metabarcode for analysing the metabarcoding data of other mock controls and commercial herbal formulations.

### 3.3. Metabarcoding of different types of mock controls

Total reads, reads obtained after filtering, and percentage of reads analysed (from filtered reads) after discarding OTU clusters having <5 reads for each mock control are shown in Supplementary Information Table S2. For the first type of control, which is gDNA pooled controls of different genus comprising 5 (5P), 10 (10P), and 15 (15P) plant species total 18657 and 452380 reads were obtained for *rbcL* and the *ITS2*, respectively (Table S2). *Zingiber officinale* had the highest percentage reads with *rbcL* metabarcode in 5P (45.9%) and 10P (30.9%), whereas *Senna tora* had the highest percentage (21.3%) in 15P (Supplementary Information Fig. S3a). *Eclipta prostrata* had the highest percentage of reads with *ITS2* metabarcode in 5P (97.02%) and 10P (79.5%), whereas *Phyllanthus emblica* had the highest percentage (40.7%) in 15P (Supplementary Information Fig. S3a). In 5P, out of five total four target plants were detected with *rbcL* metabarcode and three target plants were detected with *ITS2*. In 10P, out of ten, a total nine targeted plants were detected with *rbcL* and seven targeted plants were detected with *ITS2*. In 15P, out of fifteen, a total thirteen targeted plants were detected with *rbcL* and twelve targeted plants were detected with *ITS2* (Fig. 2; Supplementary Information Fig. S3a). On the whole, for the gDNA pooled controls of different genus, detection efficiency of *rbcL* was observed 80% for 5P and 10P, 86.7% for 15P. While, detection efficiency of *ITS2* was observed 80% and combined detection efficiency of both metabarcodes was observed 100% for all three gDNA pooled mock controls (Fig. 2a, 3). Five plant species that were present in all three gDNA pooled mock controls had 80% average fidelity and other five plant species that were present in two groups i.e., 10P and 15P had 90% average fidelity with *rbcL* metabarcode (Table 1). *ITS2* metabarcode exhibited 66.7% average fidelity for five plant species that were present in all three gDNA pooled mock controls and 90% average fidelity for other five plant species that were present in 10P and 15P. Combined average fidelity with both barcodes was 100% for gDNA pooled control (Table 1).

For the simulated plant biomass controls (i.e., blended formulations, or second type of mock controls), total 42140 and 334017 reads were obtained after filtering for *rbcL* and the *ITS2*, respectively (Table S2). The highest percentage of reads were observed for *Azadirachta indica* in 5S (62.5%) and in 10S (34.7%), whereas, in 15S *Justicia adhatoda* (24.8%) showed highest percentage of reads using *rbcL* metabarcode (Supplementary Information Fig. S3b). While in the case of *ITS2, Eclipta prostrata* had the highest percentage of reads in all simulated plant biomass controls (Supplementary Information Fig. S3b). In 5S, 10S and 15S total 3, 7, and 10 targeted plants were detected by *rbcL* and total 3, 6, and 8 targeted plants were detected by *ITS2* respectively (Fig. 2a; Supplementary Information Fig. S3b). Detection efficiency for simulated plant biomass controls was observed 80% for 5S, 70% for 10S, and 66.7% for 15S with *rbcL* metabarcode. While, detection efficiency of *ITS2* was observed 60% for all three groups of simulated plant biomass control and combined detection efficiency of both metabarcodes was observed 80% for 5S and 10S and 69.4% for 15S (Fig. 3). Five plant species that were present in all three groups of simulated plant biomass control had 80% average fidelity and other five plant species that were present in 10S and 15S had 70% average fidelity with *rbcL* metabarcode (Table 1). *ITS2* metabarcode exhibited 53.3% average fidelity for five plant species that were present in all three groups and 70% average fidelity for other five plant species that were present in two groups i.e., 10S and 15S. Combined average fidelity with both barcodes was observed 80% for simulated plant biomass controls (Table 1).

**Figure 3.**
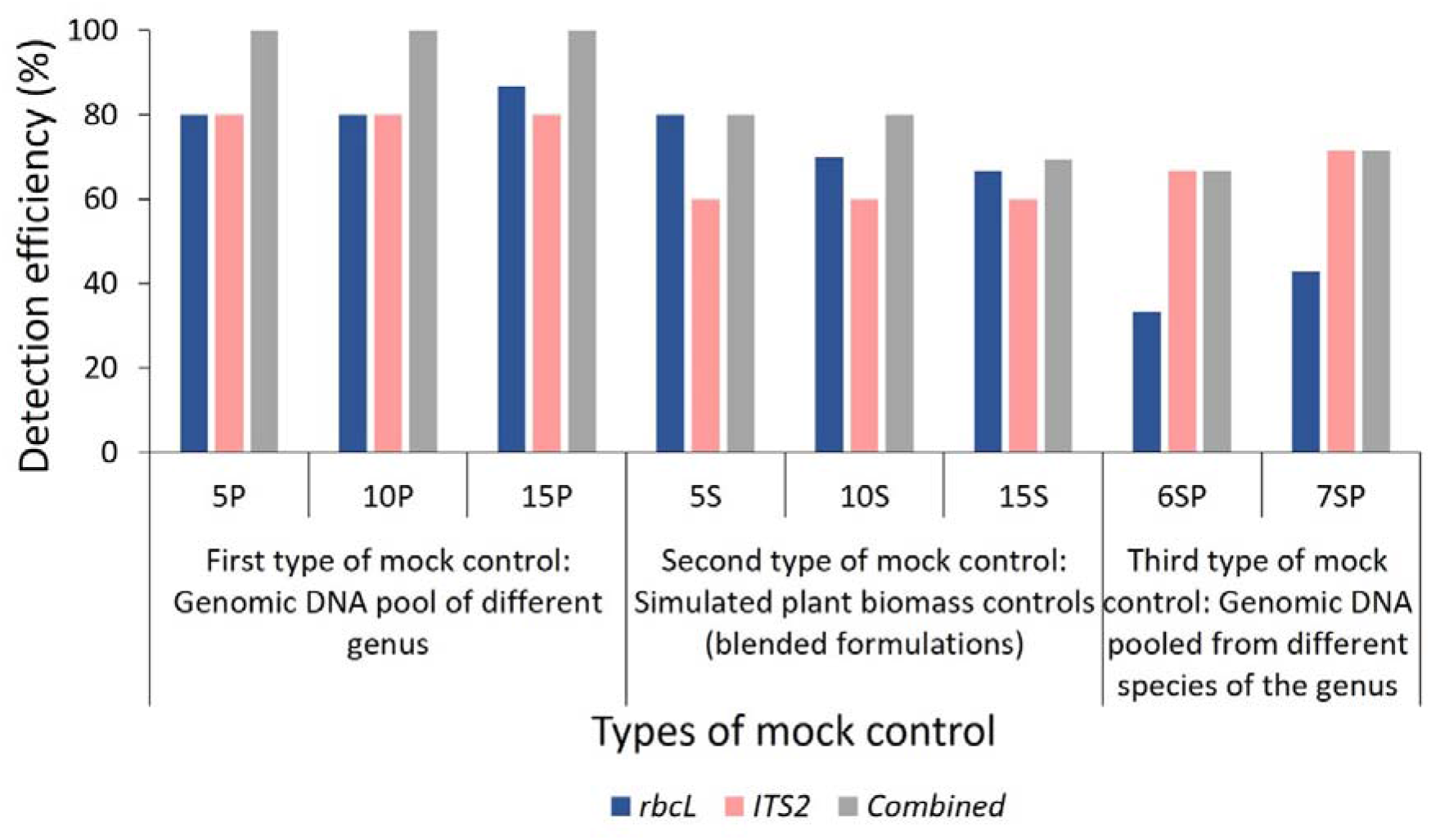
Detection efficiency obtained in mock controls by *rbcL, ITS2* and combined approach. Detection efficiency (%) was calculated using a formula described in the Metabarcoding data analysis section of Materials and Methods. 5P: genomic DNA pool of five plant species, 10P: genomic DNA pool of ten plant species, 15P: genomic DNA pool of fifteen plant species, 5S: simulated blended plant pool of five plant species, 10S: simulated blended plant pool of ten plant species, 15S: simulated blended plant pool of fifteen plant species, 6SP: genomic DNA pool of six plant species from genera *Asparagus* and *Terminalia*, 7SP: genomic DNA pool of seven plant species from genera *Piper* and *Phyllanthus*. Detailed list of plant species used in each mock control described in Fig. 2.

For the third type of control, which is gDNA pooled from different species of two genera, a total of 10265 and 164358 reads were obtained for *rbcL* and the *ITS2*, respectively (Table S2). In 6SP, out of six plant species total three plant species include *Asparagus racemosus* (81.1%), *Terminalia bellirica* (11.8%) and *Terminalia chebula* (6.8%) were detected by *rbcL* while *ITS2* metabarcode was able to resolve five plant species except *Asparagus dumosus* (Fig. 2b; Supplementary Information Fig. S3c). In 7SP, out of seven plant species two plant species include *Piper nigrum* (24.7%) *and Phyllanthus emblica* (72.3%) were detected by *rbcL*, while *ITS2* metabarcode was able to resolve four plant species include *Piper nigrum* (0.8%), *Piper longum* (21%), *Phyllanthus emblica* (62.5%), *Phyllanthus amaras* (13.5%) (Fig. 2b; Supplementary Information Fig. S3c). *rbcL* showed 33.3% and *ITS2* showed 66.7% detection efficiency in species level control with combined detection efficiency 66.7% (Fig. 2b, 2). These two species level controls indicate a greater spectrum of resolution of *ITS2* metabarcode than *rbcL* metabarcode. This finding corroborates with earlier reports in which authors demonstrated that *ITS2* metabarcode has greater species level discrimination power than *rbcL* while *rbcL* has greater universality^24,32^.

Here, in the first and third type of mock controls, DNA was pooled in equal proportions and for the second type of simulated plant biomass controls, equal weight of each plant species part that has therapeutic importance was mixed. Therefore, theoretically in metabarcoding an equal proportion of reads for each plant species should be obtained in each control. However, here the percentage of reads for each plant varied in each control and several species were not detected, showing DNA extraction biasness, primer fit compatibility and PCR amplification biases^33^. *Terminalia arjuna, T. chebula* and *Phyllanthus emblica* were not detected in simulated plant biomass control might due to variability in quality and quantity of DNA extracted from each plant species from the mixtures as different parts of plants, i.e., rhizome, fruits, leaves, and bark have been added into plant biomass controls^34^. In addition to that, *ITS2* metabarcode is unable to resolve *Andrographis paniculata* and *Justicia adhatoda* in all mock controls because of the newly designed *ITS2* metabarcode is impotent to amplify target sequence from these two plant species (Fig.1; Supplementary Information Table S1). Despite the above-mentioned limitations, the combined *rbcL* and *ITS2* metabarcoding approach was able to resolve plant species with high fidelity (Table 1) and can be implemented for the detection of plant species in herbal products.

### 3.4. Metabarcoding of single drugs

Tulsi (*Ocimum tenuiflorum*)^35^, Gokhru (*Tribulus terrestris*)^36^, Shatavari (*Asparagus racemosus*)^37^, Vasa (*Justicia adhatoda*), Ashwagandha (*Withania somnifera*), Bhringraj (*Eclipta alba*), and Arjuna (*Terminalia arjuna*) were among the 17 single drugs that were collected. Total reads, reads obtained after filtering, and percentage of analysed reads (from filtered reads) after discarding OTU clusters having <5 reads for each single drug are shown in Supplementary Information Table S3. For all single drugs, 128540 total raw reads for *rbcL* and 1211935 total raw reads for *ITS2* metabarcode were obtained (Table S3). On an average, 12.6% (0-99.8%) reads have been obtained for non-targeted plant species with *rbcL* metabarcode and 46.5% (0-100%) reads have been obtained for non-targeted plant species with *ITS2* metabarcode in single drugs (Table S4). Cross-contamination with allied species, harvesting process, pollen contamination, misidentification due to cryptic taxonomy, polynomial vernacular identification, manufacturing and packing procedure may contribute to the presence of non-targeted plant species^23^.

In tulsi powder (labeled as *Ocimum tenuiflorum*), *rbcL* was able to detect *Ocimum tenuiflorum* ranging between 99.8% to 64.4%. Along with *O. tenuiflorum*, substituted species *Ocimum basilicum*^9^ was also observed in three herbal products with 17.7, 14.4 and 1.4% reads with *rbcL* metabarcode, respectively (Fig. 4a). *O. tenuiflorum* could not be detected by *ITS2* metabarcode in any samples, however *O. basilicum* was found in one sample with 11.6% reads (Fig.4b). This could be due to inability of new *ITS2* metabarcode for amplifying *O. tenuiflorum* (Table S1) which is proof of PCR biasness toward the unintentional and low level of contamination present in samples and leads to high number of reads for non-targeted plant species.

**Figure 4.**
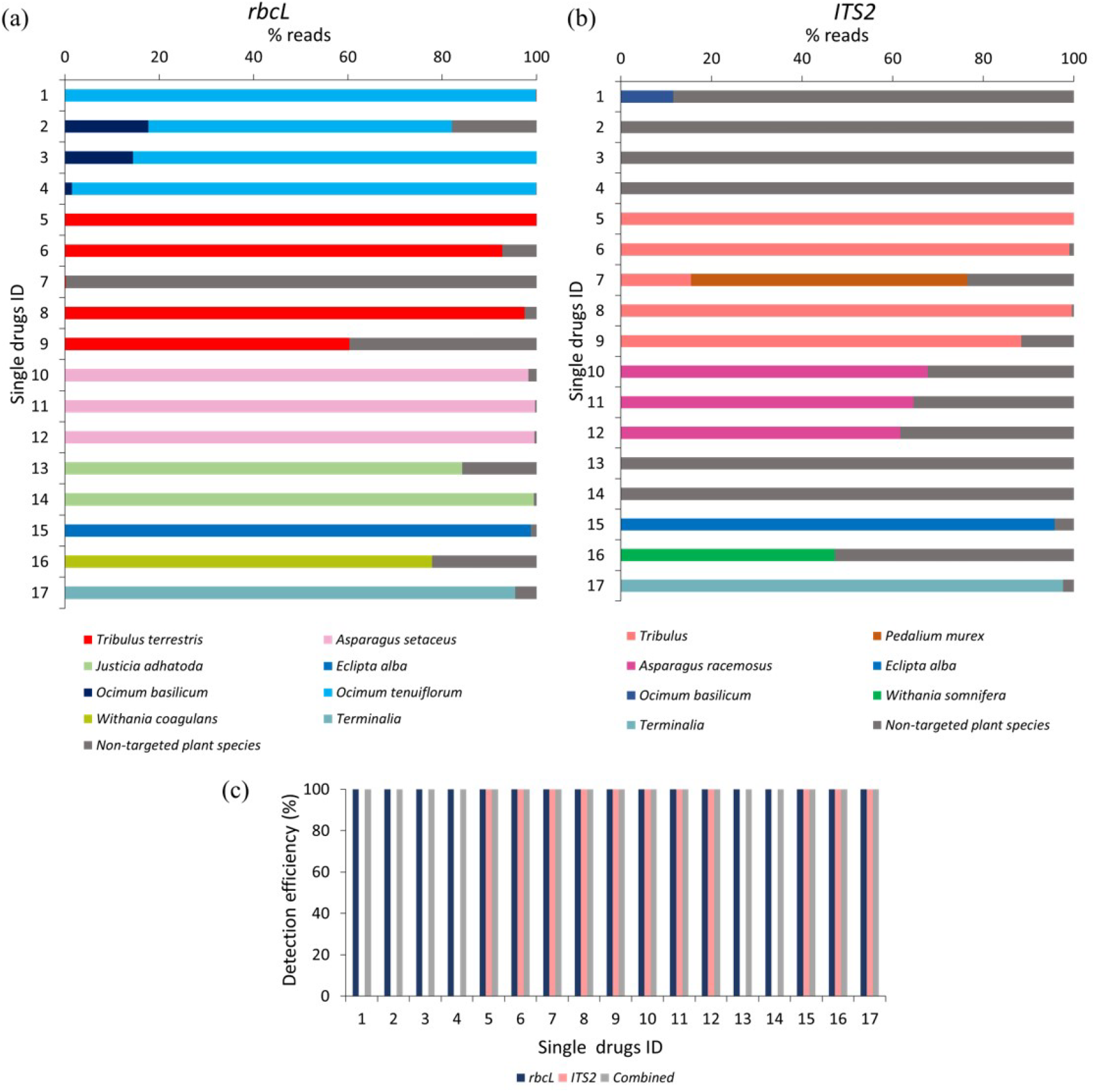
Relative abundance of the plant species and detection efficiency in single drugs through *rbcL* and *ITS2* metabarcoding. (a) Relative abundance (% reads) of the plant species detected in single drugs through *rbcL* metabarcoding. Relative abundance (% reads) of non-targeted plant species is reported in Supplementary Information Table S4. (b) Relative abundance (% reads) of the plant species detected in single drugs through *ITS2* metabarcoding. Relative abundance of non-targeted plant species is reported in Supplementary Information Table S4. (c) Detection efficiency obtained in single drugs by *rbcL, ITS2* and combined metabarcoding approach. Single drugs ID 1 to 4 for Tulsi (*Ocimum tenuiflorum*) powder, 5 to 9 for Gokhru (*Tribulus terrestris*) powder, 10 to 12 for Shatavari (*Asparagus racemosus*) powder, 13 and 14 for Vasa (*Justicia adhatoda*) powder, 15 for Bhringraj (*Eclipta alba*) powder, 16 for Ashwagandha (*Withania somnifera*) powder, and 17 for Arjuna (*Terminalia arjuna*) powder.

In Gokhru powder, *Tribulus terrestris* (Chota Gokhru) were detected up to species level by *rbcL* with 100% reads in sample 5, 92.7% in sample 6, 0.2% in sample 7, 97.4% in sample 8 and 60.3% in sample 9 (Fig. 4a). The *ITS2* metabarcode was able to resolve *T. terrestris* only at the genus level with 100% in sample 5, 99% in sample 6, 15.5% in sample 7, 99.6 % in sample 8 and 88.4% in sample 9. *ITS2* revealed the presence of 61% of *Pedalium murex* (Bada Gokhru) in sample 7 (Fig. 4b), which is a commonly substituted species and labelled as Gokhru in Indian marketed herbal products^38^.

In all three Shatavari powder, *rbcL* metabarcode detected *Asparagus setaceus* instead of *A. racemosus* with 98.3% reads in sample 10, 99.7% reads in sample 11, and 99.6% reads in sample 12. By *ITS2*, 67.7% to 61.7% reads for *A. racemosus* were obtained in all three shatavari powder. In vasa powder, *Justicia adhatoda* was detected by *rbcL* with 84.2% reads in sample 13 and 99.4% reads in sample 14. While *ITS2* was not able to resolve *Justicia adhatoda* because of the amplification inability of our *ITS2* metabarcode (Fig. 4, Supplementary Information Table S1). This result was consistent with the mock control 15P and 15S. In Ashwagandha powder (sample 16), *Withania coagulans* (77.9% reads) instead of *W. somnifera* was detected by *rbcL*. While with *ITS2, W. somnifera* was detected with 47.2% reads.

In Bhringraj powder, both *rbcL* and *ITS2* metabarcode were able to detect *Eclipta alba* with 98.8% and 95.7% reads respectively. In Arjuna powder, *rbcL* metabarcode was able to identify *Terminalia arjuna* only up to genus level with 95.5% reads while *ITS2* metabarcode was identified *T. arjuna* at species level with 97.6% reads (Fig. 4).

For Tulsi, Gokhru, Shatavari, and Vasa powder 100% fidelity was obtained with *rbcL* metabarcode, while *ITS2* metabarcode exhibited 0% fidelity for Tulsi and Vasa powder and 100% fidelity for Gokhru and Shatavari powder (Table 2a). Suggesting that, for authentication of Tulsi and Vasa powder our *rbcL* metabarcode works efficiently but not *ITS2*. While, *rbcL* metabarcode was not suitable for identification of *Pedalium murex* in Gokhru powder as *rbcL* sequence for *Pedalium murex* was not available in NCBI (database was accessed on December 15, 2022). In addition, *Terminalia arjuna* was resolved only up to genus level by *rbcL* and *Terminalia terrestris* was resolved up to only at the genus level by *ITS2* metabarcode. This is in concordance with mock controls and could be due to low interspecific variability of barcode sequence covered by our metabarcodes for resolving species to be identified. Although, combined metabarcoding approach provides 100% detection efficiency with 100% fidelity for single drugs by overcoming the limitations of individual barcodes due to PCR biases, low interspecific variability or the absence of the corresponding sequences in the database. Seethapathy et al.^23^ demonstrated 67% fidelity for targeted plant species present in single drugs of the European market using *ITS1* and *ITS2* barcodes. Here, we obtained 100% fidelity for targeted plant species within single drugs. The overall results of the single drugs revealed that a multi-barcode metabarcoding approach can be used to assess the prevalence of widespread adulterated and substituted plant material in single drugs, as well as the implementation of more stringent supply chain precautionary measures at primary level.

### 3.5. Metabarcoding of polyherbal formulations

Trikatu, Sitopaladi, Rasayana, Hingwashtak and Talisadi (Talisadya)^39^ were among the 15 polyherbal formulations that were collected. Total reads, reads obtained after filtering, and percentage of analysed reads (from filtered reads) after discarding OTU clusters having <5 reads for each polyherbal formulation are shown in Supplementary Information Table S5. A total of 53087 and 1429238 reads were obtained by *rbcL* and *ITS2* metabarcode for polyherbal formulation respectively (Table S5). On average, 1.4% (0-3.9%) reads have been obtained for non-targeted plant species with *rbcL* metabarcode and 16.5% (0.1-87.4%) reads have been obtained for non-targeted plant species with *ITS2* metabarcode in polyherbal formulations (Table S6).

In Trikatu, 28.2%, 4.6%, and 57% reads for *Piper nigrum* with *rbcL* and 8.3%, 14.7 and 37.9% reads with *ITS2* was observed in sample 18, 19, and 20 respectively. *Piper longum* comprised 2%, 37.1%, and 47.8% reads with *rbcL* and 0%, 42.1% and 19.7% with *ITS2*; *Zingiber officinale* possess 67.6%, 56.5%, and 43.6% with *rbcL* and 4.3 %, 18.8 % and 2.4 % with *ITS2* in sample 18, 19, and 20 respectively (Fig. 5a, 5b). All three targeted plants were detected (i.e., 100% detection efficiency) in all three Trikatu samples (i.e., 100% fidelity) using a combined approach (Fig. 5c, Table 2b). Nevertheless, *ITS2* showed the higher percentage of non-targeted reads (87.4% in sample 18, 24.4% in sample 19, and 40% in sample 20) might be due to technical bias that can be introduced during DNA extraction and PCR towards the unintentional cross contamination happens during the supply chain (Fig. 5b).

**Figure 5.**
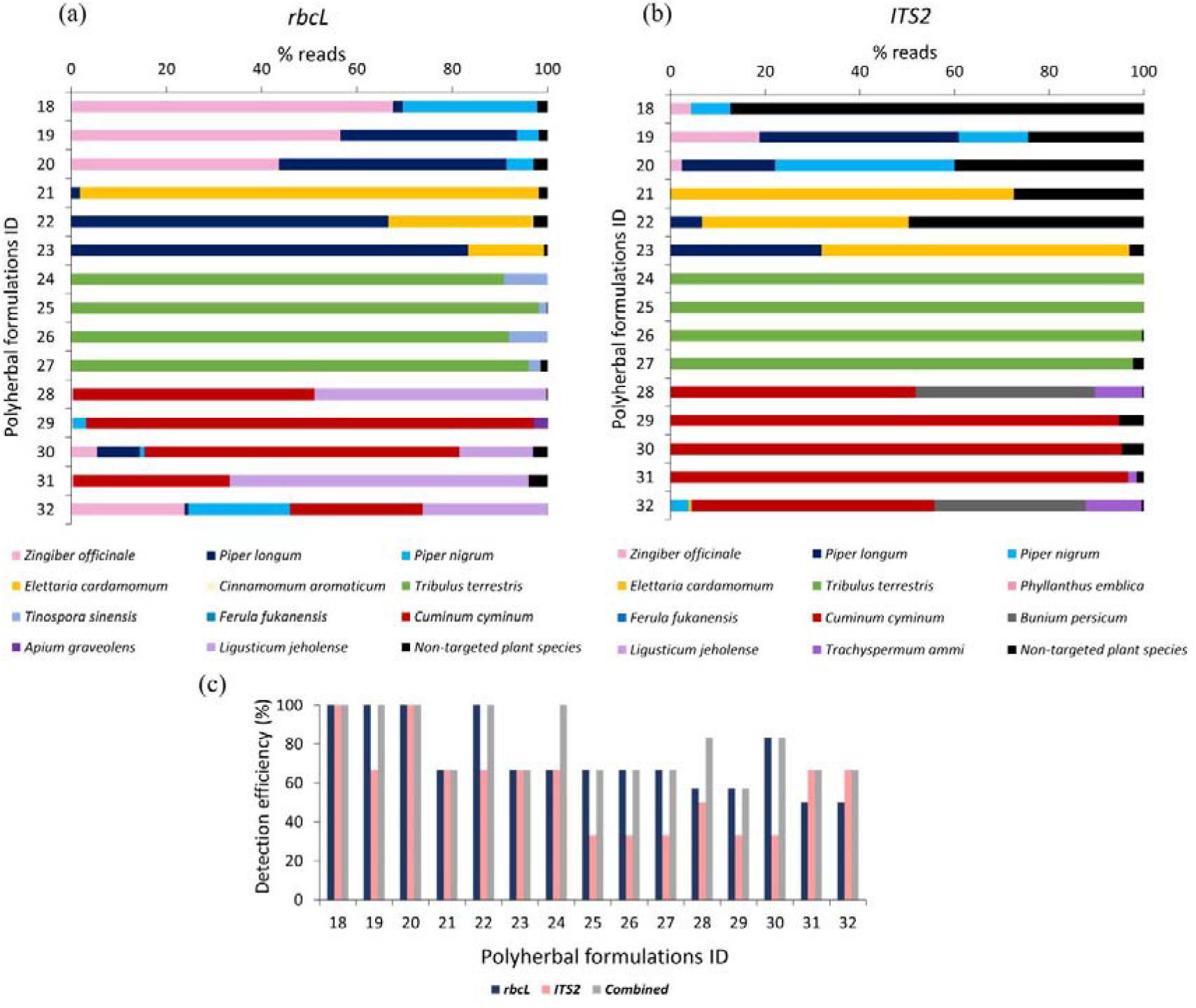
Relative abundance of the plant species and detection efficiency in polyherbal formulations through *rbcL* and *ITS2* metabarcoding. (a) Relative abundance (% reads) of the plant species detected in polyherbal formulations through *rbcL* metabarcoding. Relative abundance (% reads) of non-targeted plant species is reported in Supplementary Information Table S6. (b) Relative abundance (% reads) of the plant species detected in polyherbal formulations through *ITS2* metabarcoding. Relative abundance (% reads) of non-targeted plant species is reported in Supplementary Information Table S6. (c) Detection efficiency obtained in polyherbal formulation by *rbcL, ITS2* and combined metabarcoding approach. Polyherbal formulations ID 18 to 20 for Trikatu powder, 21 to 23 for Sitopaladi powder, 24 to 27 for Rasayana powder, 28 to 31 for Hingwashtak powder, 32 for Talisadi powder.

Sitopaladi powder is primarily composed of five constituents; with the exclusion of sacharium (sugar) and vanshlochan (the female bamboo exudate) and from the total number of designated species, the aim was to detect *Cinnamomum cassia, Piper longum*, and *Elettaria cardamom. Cinnamomum cassia* was detected by *rbcL* metabarcode with 0.2% reads only in sample 22. *Piper longum* exhibited 1.8, 66.6, and 83.3% of reads with *rbcL* and 0.1, 6.6, and 31.9% reads with *ITS2* in samples 21, 22, and 23 respectively. *Elettaria cardamomum* showed 96.3%, 30.2%, and 15.9% reads with *rbcL* and 72.4, 43.7 and 65.1% with *ITS2* in samples 21, 22, and 23 respectively (Fig. 5a, 5b). Overall, the combined approach showed 66.7% detection efficiency in sample 21, 100% in sample 22, and 66.7% in sample 23 and average 77.8% fidelity (Fig. 5c, Table 2b).

In Rasayana, *rbcL* metabarcode exhibited 91% to 98 % reads for *Tribulus teresteris* while *ITS2* metabarcode exhibited 98 to 100% reads in all Rasayana samples (sample 24 to 27). *Tinospora cordifolia* was resolved by *rbcL* in all samples with 1.7% to 9.1% reads while *ITS2* metabarcode could not. *rbcL* was able to resolve *Tinospora cordifolia* in all samples with percentage of reads ranging from 1.7 to 9.1, while *ITS2* metabarcode could not (Fig. 5a, 5b). *Phyllanthus emblica* was not detected by both metbarcode except in sample 24 (0.01% reads were obtained by *ITS2*), possibly due to DNA extraction biasness as DNA extraction from amla fruits is difficult due to high acidic nature and high tannin content^40^. The combined metabarcoding approach showed 100% detection efficiency in sample 24 and 66.7% in remaining other samples with average 75% fidelity (Fig. 5c, Table 2b).

Hingwashtak powder (sample 28 to 31) comprised seven ingredients include *Zinger officinale, Piper nigrum, Piper longum, Cyminum cyminum, Carum carvi* [*Cyminum cyminum and Carum carvi* commonly substituted with *Bunium persicum* (Syn. *Elwendia persica*)^41–43^], *Apium leptophyllum* (majority of commercial products comprised/labelled *Apium graveolens* instead of *Apium leptophyllum*, further these plant species commonly substituted with *Trachyspermum ammi*^44^) and *Ferula foetida*. From these seven ingredients, *rbcL* metabarcode was able to resolve *Zinger officinale* in all samples with 0.37 to 5.5% reads, *Piper nigrum* in sample 29 (2.7% reads) and 30 (1.0% reads), *Piper longum* in sample 30 (8.9% reads), *Cyminum cyminum* in all samples with 32.7% to 94.1% reads, *Apium graveolens* in sample 29 (2.9% reads). *Carum carvi* commonly substituted with *Bunium persicum* (*Elwendia persica*), neither *Carum carvi* nor *Bunium persicum* was detected by *rbcL* in all Hingwashtak samples (Fig. 5a). *ITS2* metabarcode exhibited high prevalence of *Cyminum cyminum* in all samples with 51.8% to 95.4% reads, *Bunium persicum* in sample 28 with 37.9% reads, *Trachyspermum ammi* (substitution of *Apium leptophyllum* or *Apium graveolens*) in sample 28 with 10% reads (Fig. 5b). In addition, *ITS2* metabarcode was able to detect *Trachyspermum ammi* also in sample 29 and 31. *RbcL* metabarcode showed reads for *Ligusticum jeholense* (Chinese medicinal herb from the Apiaceae family) instead of *Trachyspermum ammi* in sample 28, 30, 31 and *Apium graveolens* in sample 29. This could be due to our *rbcL* metabarcode unable to resolve *Trachyspermum ammi* and detect *Ligusticum jeholense* falling under the same family. Overall, the combined metabarcoding approach showed average 72.6% detection efficiency with 63.3% fidelity for Hingwashtak powders (Fig. 5c, Table 2b).

Talisadi/Talisadya powder (sample 32) comprises eight constituents include *Abies webbiana, Piper longum, Piper nigrum, Zinger officinale, Elettaria cardamomum, Cinnamomum Zeylanicum*, Vanshlochan (the female bamboo exudate), and sugar (sugar and Vanshlochan are excluded for metabarcoding analysis). From these six ingredients, only 3 plant species which include *Piper longum* (0.8% reads), *Piper nigrum* (21.3% reads), and *Zinger officinale* (23.8% reads) were detected by *rbcL* metabarcode. Three plant species which include *Piper nigrum* (3.79 % reads), *Zinger officinale* (0.03% reads) and *Elettaria cardamomum* (0.6% reads) were detected by *ITS2* metabarcode. *Abies webbiana* and *Cinnamomum Zeylanicum* were not detected by either of the barcodes. The combined metabarcode approach was able to detect four plant species (66.7% detection efficiency) out of six plant species (Fig. 5c). In Talisadi/Talisadya powder, *Cuminum cyminum* (27.9% reads with *rbcL* and 51.3% reads with *ITS2*), *Bunium persicum* (32.0% reads with *ITS2*), *Ligusticum jeholense* (26.2% reads with *rbcL*) and *Trachyspermum ammi* (11.8% reads with *ITS2*) was detected might be due to unintentional cross contamination happen during sample processing as collection of Hingwashtak powder sample 28 and Talisadi powder sample 32 were done from the same company. In addition, a high percentage of reads were covered by *Cuminum cyminum, Trachyspermum ammi, Ligusticum jeholen*s, *Bunium persicum* (all plant species belongs to Apiaceae family), then *Zingiber officinale, Piper longum*, and *Piper nigrum*. This could be because of technical bias that can be introduced during DNA extraction and PCR.

### 3.6. Fidelity of targeted plant species

Up to this point, the fidelity of plant species per number of mock controls or herbal formulations has been calculated and discussed. Here, we have estimated the fidelity of the targeted plant species included within different mock controls as well as within different polyherbal formulations to get better perspective of species discrimination capabilities and reliabilities of single and multi-barcode approach. Both the rbcL and ITS2 metabarcodes were able to resolve 19 (46.7%) of the 39 listed plant species at the species level. However, plants detected at the species level were different, and a multi-barcode approach provided species-level resolution for 27 (69.23%) species, leading to a 20.5% increment in the whole (Table 3). This confirms robustness for our newly designed metabarcodes in detecting plant species at lower taxonomic level. In addition, 100% fidelity was observed for *Terminalia bellirica, Andrographis paniculata, Azadirachcta indica, Eclipta alba*, and *Centella asiatica* within gDNA controls, biomass controls, and cumulative analysis by the combined approach of *rbcL* and *ITS2*. However, the combined approach of *rbcL* and *ITS2* exhibited 100% fidelity for *Zingiber officinale, Vitex negundo, Piper nigrum, Aegale marmelous, Terminalia arjuna*, and *Phyllanthus embilica* only within gDNA controls and showed lower fidelity in biomass controls and cumulative analysis (Table 3). This could be due to biases in the DNA isolation process; yielding equal proportional DNA from the poly formulation is not possible due to genome size differences as well as differences in plant parts and amounts of secondary metabolites. Furthermore, because herbal products are intensively processed, the extracted DNA is degraded. The PCR conditions and reactions will also have a significant impact on the primer fit and PCR bias of the mixture. This was clearly demonstrated by comparing the combined fidelity of gDNA controls, biomass controls, and cumulative analysis (Table 3). On average, 83.6% fidelity was observed for targeted plant species in the cumulative analysis. This result confirmed the high reliability of our multi-barcode sequencing approach.

## 4. Conclusion

On the whole, our findings suggest that the multi-barcode DNA metabarcoding method assessed in this study can provide a composition of more diverse sets of single drugs and polyherbal formulations listed in the Ayurvedic Pharmacopoeia of India. Through the multibarcode sequencing approach, we obtained 100% an average detection efficiency and relative fidelity of targeted plants for single drugs and 79% for polyherbal formulations. We have primarily focused on detected plant species in herbal products rather than undetected plant species because there are many steps, such as DNA extraction biasness, PCR biases, and manufacturing processes, that can lead to DNA degradation or loss beyond detectable limits, resulting in failure to detect plant species. The presence of non-targeted plant species in herbal products could be due to unintentional contamination of the supply chain, economically motivated adulteration, and/or admixture of other species. Our study showed that the *rbcL* metabarcode had better detection ability for certain plant species, e.g., *Ocimum tenuiflorum, Justicia adhatoda*, and *Andrographis paniculata*, while *ITS2* had better discrimination power for certain plant species, e.g., species of the genus *Terminalia, Asparagus, Piper, Phyllanthus*, and *Pedalium murex*. Thus, the complementary approach of both metabarcodes is a promising tool for quality evaluation of herbal products and pharmacovigilance. However, the development of standardised methods for metabarcoding sequencing and bioinformatics analysis pipeline and curated database is needed for effective use as a regulatory tool to authenticate herbal products in combination with advanced chemical methods that are used to identify bioactive therapeutics.

## Funding

Gujarat State Biotechnology Mission (GSBTM), Gandhinagar, Gujarat, India, has provided financial support for the project under the Research Support Scheme, grant ID GSBTM/JDRD/584/2018/204.

## Acknowledgment

The authors would like to thank Prof. Padamnabhi S. Nagar, The Maharaja Sayajirao University of Baroda, Gujarat, India and Director, DMAPR, Gujarat, India for helping us in plant collection and authentication. The authors would like to thank Dr. Darshan Parmar, M.D. (Rasashastra and Bhaishajya Kalpana), Government Ayurvedic College, Vadodara (Gujarat, India) for providing some of the herbal formulations and Mr. Nitin Savaliya, Technical Assistant from Thermo Fisher Scientific, for NGS instrument handling and run setup.

## Conflict of Interest

Authors have no conflict of interest.

## Author contribution

TT: Performed experiments, established metabarcoding data analysis pipeline, Data analysis, Writing and Editing manuscript, Validation of final manuscript; APS: Performed experiments, Data analysis, Writing and Editing manuscript, validation of final manuscript; RP: Designed primers for metabarcoding, Performed experiments, established metabarcoding data analysis pipeline, Manuscript editing; SS: Performed experiments, Manuscript editing; CJ: Project administration, Methodology, Supervision, and Review & Editing; MJ: Principal Investigator, Conceptualization, Methodology, Supervision, and Review & Editing.

## Notes

### Competing Interest Statement

The authors have declared no competing interest.

