## Supplementary material for "DNA metabarcoding using new *rbcL* and *ITS2* metabarcodes collectively enhance detection efficiency of medicinal plants in single and polyherbal formulations": Table 1

**Table 1** Detection fidelity in mock controls^≠^.

| **Sr. no.** | **Plant Species** | **gDNA pooled mock controls** | | | | **Simulated plant biomass controls (Blended formulations)** | | | |
| --- | --- | --- | --- | --- | --- | --- | --- | --- | --- |
|  |  |  | **Relative fidelity/species (%)** | | |  | **Relative Fidelity/species (%)** | | |
|  |  | **Species present in no. of mock controls^#^** | ***rbcL*** | ***ITS2*** | **Combined** | **Species present in no. of mock controls^®^** | ***rbcL*** | ***ITS2*** | **Combined** |
| 1 | *Andrographis paniculata* | 3 | 100 | 0 | 100 | 3 | 100 | 0 | 100 |
| 2 | *Azadirachcta indica* |  | 100 | 33.3 | 100 |  | 100 | 66.7 | 100 |
| 3 | *Eclipta alba* |  | 100 | 100 | 100 |  | 100 | 100 | 100 |
| 4 | *Piper nigrum* |  | 0 | 100 | 100 |  | 100 | 100 | 100 |
| 5 | *Zingiber officinale* |  | 100 | 100 | 100 |  | 0 | 0 | 0 |
| **Average Fidelity (%)** | | | **80** | **66.7** | **100** |  | **80** | **53.3** | **80** |
| 6 | *Aegale marmelous* | 2 | 100 | 50 | 100 | 2 | 100 | 50 | 100 |
| 7 | *Centella asiatica* |  | 100 | 100 | 100 |  | 100 | 100 | 100 |
| 8 | *Terminalia arjuna* |  | 50 | 100 | 100 |  | 0 | 0 | 0 |
| 9 | *Terminalia bellerica* |  | 100 | 100 | 100 |  | 50 | 100 | 100 |
| 10 | *Vitex negundo* |  | 100 | 100 | 100 |  | 100 | 100 | 100 |
| **Average Fidelity (%)** | | | **90** | **90** | **100** |  | **70** | **70** | **80** |

^#^3: Species present in all three groups of gDNA pooled mock control (n=3); 2: Species present in two groups of gDNA pooled mock control i.e., 5P and 10P (n=2). ^®^3: Species present in all three groups of simulated plant biomass mock control (n=3); 2: Species present in two groups of simulated plant biomass mock control i.e., 5S and 10S (n=2). ^#®^Fidelity was not calculated for five plant species that present only in 15P and15S (n=1). **^≠^** Fidelity for the third type of mock control (i.e., gDNA pooled from different species of the genus) was not calculated as each plant species present only in one group (n=1).

**Table 2** Detection fidelity of single drugs and polyherbal formulations.

**Table 2a** Detection fidelity of single drugs.

| **Name of Herbal product** | **Composition of formulations** | **No. of products** | **Absolute fidelity**^#^ **/single drug** | | | **Relative fidelity (%)/single drug** | | |
| --- | --- | --- | --- | --- | --- | --- | --- | --- |
|  |  |  | ***rbcL*** | ***ITS2*** | **Combined** | ***rbcL*** | ***ITS2*** | **Combined** |
| Tulsi | *Ocimum tenuiflorum* | 4 | 4 | 0 | 4 | 100 | 0 | 100 |
| Gokhru | *Tribulus terrestris* | 5 | 5 | 5 | 5 | 100 | 100 | 100 |
| Shatavari | *Asparagus racemosus* | 3 | 3 | 3 | 3 | 100 | 100 | 100 |
| Vasa | *Justicia adhatoda* | 2 | 2 | 0 | 2 | 100 | 0 | 100 |
| **Cumulative fidelity of single drugs (%)** | | | | | | **100** | **50** | **100** |

^#^Fidelity was not calculated for Bhurgraj, Ashawgandha and Arjuna where n=1.

**Table 2b** Detection fidelity of polyherbal formulations.

| **Name of Herbal product** | **Composition of formulations** | **No. of products** | **Absolute fidelity of detection/species/polyherbal formulation^≠^** | | | **Relative fidelity of detection/ species/polyherbal formulation** | | | **Average relative fidelity (%)/ polyherbal formulations** | | |
| --- | --- | --- | --- | --- | --- | --- | --- | --- | --- | --- | --- |
|  |  |  | ***rbcL*** | ***ITS2*** | **Combined** | ***rbcL*** | ***ITS2*** | **Combined** | ***rbcL*** | ***ITS2*** | **Combined** |
| Trikatu | *Zinger officinale* | 3 | 3 | 3 | 3 | 100 | 100 | 100 | 88.9 | 100 | 100 |
|  | *Piper nigrum* |  | 3 | 3 | 3 | 100 | 100 | 100 |  |  |  |
|  | *Piper longum* |  | 2 | 3 | 3 | 66.7 | 100 | 100 |  |  |  |
| Sitopaladi | *Piper logum* | 3 | 3 | 3 | 3 | 100 | 100 | 100 | 77.8 | 66.7 | 77.8 |
|  | *Eletaria cardamom* |  | 3 | 3 | 3 | 100 | 100 | 100 |  |  |  |
|  | *Cinnamomum cassia* |  | 1 | 0 | 1 | 33.3 | 0 | 33.3 |  |  |  |
| Rasayana | *Tribulus terrestris* | 4 | 4 | 4 | 4 | 100 | 100 | 100 | 75 | 33.3 | 75 |
|  | *Tinospora sinensis* |  | 4 | 0 | 4 | 100 | 0 | 100 |  |  |  |
|  | *Phyllanthus emblica* |  | 1 | 0 | 1 | 25 | 0 | 25 |  |  |  |
| Hingwashtak | *Zinger officinale* | 4 | 4 | 1 | 4 | 100 | 25 | 100 | 57.1 | 39.3 | 63.3 |
|  | *Piper nigrum* |  | 2 | 0 | 2 | 50 | 0 | 50 |  |  |  |
|  | *Piper longum* |  | 1 | 2 | 2 | 25 | 50 | 50 |  |  |  |
|  | *Apium graveolens* (substituted with *Trachyspermum ammi*) |  | 4 | 3 | 4 | 100 | 75 | 100 |  |  |  |
|  | *Cyminum cyminum* |  | 4 | 4 | 4 | 100 | 100 | 100 |  |  |  |
|  | *Carum carvi* [substituted with *Bunium persicum (Elwendia persica)]* |  | 0 | 1 | 1 | 0 | 25 | 25 |  |  |  |
|  | *Ferula foetida* |  | 1 | 0 | 1 | 25 | 0 | 25 |  |  |  |

^≠^Fidelity was not calculated for Talisadi/Talisadya as n=1.

**Table 3** Fidelity of targeted plant species present within mock controls as well herbal formulations.

| Plant species | Family | Plant part used in simulated biomass mock control or formulations | Resolution at taxa level | | A number of gDNA mock controls in which plant species present | Relative fidelity of each plant species presents within first and third type of mock controls (gDNA controls) | | | A total number of biomass controls, single drugs and polyherbal formulations in which plant species present | Relative fidelity of each plant species presents within second type of mock control (simulated biomass control), single drugs and polyherbal formulations (biomass controls + herbal formulations) | | | A number of mock controls and herbal products in which plant species present | Relative fidelity of each plant species presents in different types of mock controls and herbal products (cumulative analysis) | | |
| --- | --- | --- | --- | --- | --- | --- | --- | --- | --- | --- | --- | --- | --- | --- | --- | --- |
|  |  |  | ***rbcL*** | ***ITS2*** |  | ***rbcL*** | ***ITS2*** | **Combined** |  | ***rbcL*** | ***ITS2*** | **Combined** |  | ***rbcL*** | ***ITS2*** | **combined** |
| *Andrographis paniculata* | Acanthaceae | Whole plant | Species | ND | 3 | 100 | 0 | 100 | 3 | 100.0 | 0.0 | 100.0 | 6 | 100.0 | 0.0 | 100.0 |
| *Azadirachcta indica* | Meliaceae | Leaves | Species | ND | 3 | 100 | 33.3 | 100 | 3 | 100.0 | 66.7 | 100.0 | 6 | 100.0 | 50.0 | 100.0 |
| *Eclipta alba* | Asteraceae | Whole plant | Species | Species | 3 | 100 | 100 | 100 | 4 | 100.0 | 100.0 | 100.0 | 7 | 100.0 | 100.0 | 100.0 |
| *Piper nigrum* | Piperaceae | Seed | Species | Species | 4 | 100 | 100 | 100 | 11 | 81.8 | 90.9 | 90.9 | 15 | 86.7 | 93.3 | 93.3 |
| *Zingiber officinale* | Zingiberaceae | Rhizome | Family | Family | 3 | 100 | 100 | 100 | 11 | 54.5 | 63.6 | 63.6 | 14 | 64.3 | 71.4 | 71.4 |
| *Aegale marmelous* | Rutaceae | Fruits | Family | ND | 2 | 100 | 50 | 100 | 2 | 50.0 | 0.0 | 50.0 | 4 | 75.0 | 25.0 | 75.0 |
| *Centella asiatica* | Apiaceae | Whole plant | Species | Species | 2 | 100 | 100 | 100 | 2 | 100.0 | 100.0 | 100.0 | 4 | 100.0 | 100.0 | 100.0 |
| *Terminalia arjuna* | Combretaceae | Bark | Genus | Species | 3 | 0 | 100 | 100 | 3 | 33.3 | 33.3 | 33.3 | 6 | 16.7 | 66.7 | 66.7 |
| *Terminalia bellerica* | Combretaceae | Fruits | Species | Species | 3 | 100 | 100 | 100 | 2 | 0.0 | 100.0 | 100.0 | 5 | 60.0 | 100.0 | 100.0 |
| *Vitex negundo* | Lamiaceae | Leaves | Species | species | 2 | 100 | 100 | 100 | 2 | 50.0 | 50.0 | 50.0 | 4 | 75.0 | 75.0 | 75.0 |
| *Bacopa monneri* | Plantaginaceae | Whole plant | Species | Species | 1 | NA | NA | NA | 1 | NA | NA | NA | 2 | 50.0 | 100.0 | 100.0 |
| *Cassia tora* | Fabaceae | Leaves | Species | Species | 1 | NA | NA | NA | 1 | NA | NA | NA | 2 | 50.0 | 100.0 | 100.0 |
| *Phyllanthus embilica* | Phyllanthaceae | Fruits | Species | Species | 2 | 100 | 100 | 100 | 5 | 0.0 | 20.0 | 20.0 | 7 | 28.6 | 42.9 | 42.9 |
| *Terminalia chebula* | Combretaceae | Fruits | Genus | Species | 2 | 100 | 100 | 100 | 1 | NA | NA | NA | 3 | 66.7 | 66.7 | 100.0 |
| *Justicia adhatoda* | Acanthaceae | Leaves | Species | ND | 1 | NA | NA | NA | 3 | 100.0 | 0.0 | 100.0 | 4 | 100.0 | 0.0 | 100.0 |
| *Piper longum* | Piperaceae | Fruits | Species | Species | 1 | NA | NA | NA | 11 | 63.6 | 72.7 | 72.7 | 12 | 58.3 | 75.0 | 75.0 |
| *Asparagua racemosus* | Asparagaceae | Root | Genus | Species | 1 | NA | NA | NA | 3 | 100.0 | 100.0 | 100.0 | 4 | 100.0 | 100.0 | 100.0 |
| *Tribulus terrestris/*  *Pedalium murex* | Zygophyllaceae/  Pedaliaceae | Fruits | Species/  ND | Genus/  Species |  |  |  |  | 9 | 100.0 | 100.0 | 100.0 | 9 | 100.0 | 100.0 | 100.0 |
| *Ocimum tenuiflorum/*  *Ocimum basilicum* | Lamiaceae | Leaves | Species/  Species | ND/ND |  |  |  |  | 4 | 100.0 | 0.0 | 100.0 | 4 | 100.0 | 0.0 | 100.0 |
| *Elettaria cardamomum* | Zingiberaceae | Seed | Species | Species |  |  |  |  | 4 | 75.0 | 100.0 | 100.0 | 4 | 75.0 | 100.0 | 100.0 |
| *Carum carvi/*  *Elwendia persica* | Apiaceae | Seed | ND/ND | ND/  Genus |  |  |  |  | 4 | 0.0 | 25.0 | 25.0 | 4 | 0.0 | 25.0 | 25.0 |
| *Cinnamomum cassia* | Lauraceae | Bark | Genus | ND |  |  |  |  | 4 | 25.0 | 0.0 | 25.0 | 4 | 25.0 | 0.0 | 25.0 |
| *Cyminum cyminum* | Apiaceae | Seed | Species | Species |  |  |  |  | 4 | 100.0 | 100.0 | 100.0 | 4 | 100.0 | 100.0 | 100.0 |
| *Ferula foetida* | Apiaceae | Gum resin | ND | Genus |  |  |  |  | 4 | 0.0 | 25.0 | 25.0 | 4 | 0.0 | 25.0 | 25.0 |
| *Tinospora sinensis* | Menispermaceae | Root | Species | ND |  |  |  |  | 4 | 100.0 | 0.0 | 100.0 | 4 | 100.0 | 0.0 | 100.0 |
| *Apium leptophyllum*  */Trachyspermum ammi/Apium graveolens* | Apiaceae | Seed | ND/ND/Species | ND/  Species/ND |  |  |  |  | 4 | 100.0 | 75.0 | 100.0 | 4 | 100.0 | 75.0 | 100.0 |
| *Withania somnifera* | Solanaceae | Root | Genus | Species |  |  |  |  | 1 | NA | NA | NA | 1 | NA | NA | NA |
| *Abies webbiana* | Pinaceae | Leaves | ND | ND |  |  |  |  | 1 | NA | NA | NA | 1 | NA | NA | NA |
| *Asparagua dumosus* | Asparagaceae | Leaves | ND | ND | 1 | NA | NA | NA |  |  |  |  | 1 | NA | NA | NA |
| *Asparagus adscendens* | Asparagaceae | Leaves | ND | Species | 1 | NA | NA | NA |  |  |  |  | 1 | NA | NA | NA |
| *Phyllanthus amaras* | Phyllanthaceae | Leaves | ND | Species | 1 | NA | NA | NA |  |  |  |  | 1 | NA | NA | NA |
| *Phyllanthus madresapatensis* | Phyllanthaceae | Leaves | ND | ND | 1 | NA | NA | NA |  |  |  |  | 1 | NA | NA | NA |
| *Piper betle* | Piperaceae | Leaves | ND | ND | 1 | NA | NA | NA |  |  |  |  | 1 | NA | NA | NA |
| *Pipper arabarum* | Piperaceae | Leaves | ND | ND | 1 | NA | NA | NA |  |  |  |  | 1 | NA | NA | NA |

ND: Not detected, NA: Species not applicable for fidelity calculation as present only either in one mock control or either in one herbal formulations (n=1). Plant species represent in red colour are substituted plant species.
