## Supplementary Information Fig. S1, Supplementary Information Table S1 for "DNA metabarcoding using new *rbcL* and *ITS2* metabarcodes collectively enhance detection efficiency of medicinal plants in single and polyherbal formulations"

**Figure S1**

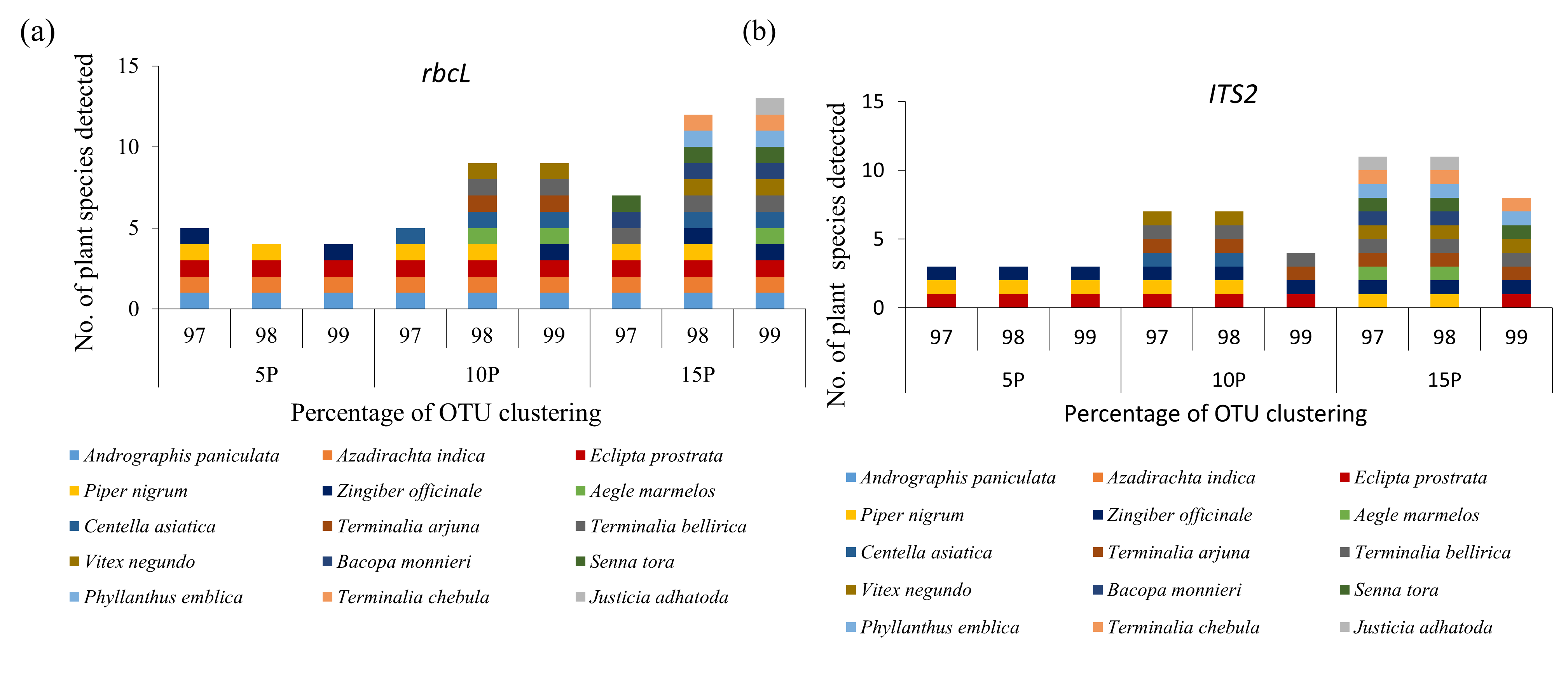

**Figure S1** The distribution of predefined herbal species detected within gDNA pool mock controls (first type of mock controls) using *ITS2* and *rbcL* metabarcode with 97%, 98%, and 99% OTU clustering.

**Figure S2**

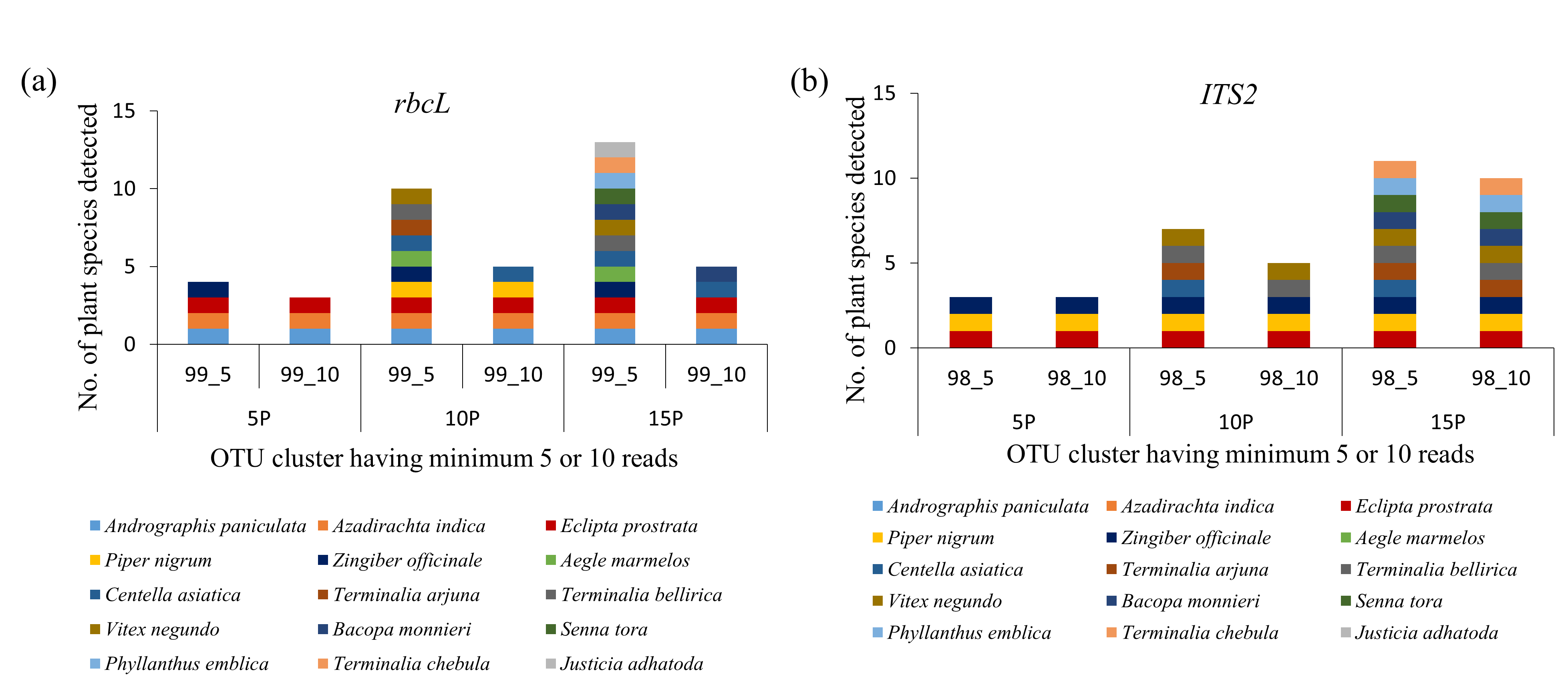

**Figure S2** The distribution of predefined herbal species detected within gDNA pool mock controls (first type of mock controls) using *ITS2* and *rbcL* metabarcode with clustered having <5 and <10 reads were discarded.

**Figure S3**

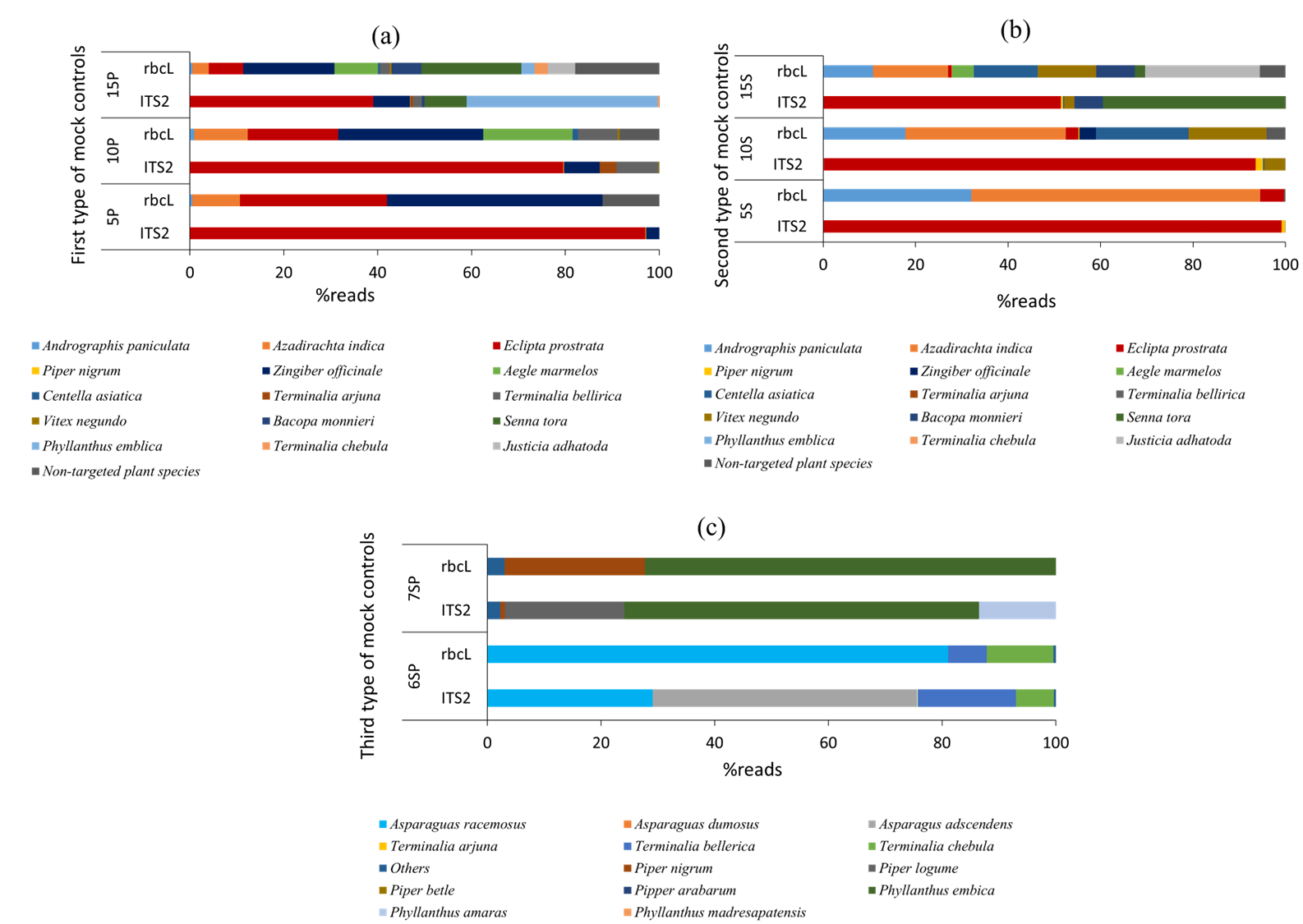

**Figure S3** Relative abundance of the plant species detected in each type of mock control through *rbcL* and *ITS2* metabarcoding sequencing. (a) Relative abundance (% reads) of the plant species detected in the first type of mock controls (i.e., genomic DNA pools of different genus) through *rbcL* and *ITS2* metabarcoding. (b) Relative abundance (% reads) of the plant species detected in the second type of mock controls (i.e., simulated plant biomass control or blended formulations) through *rbcL* and *ITS2* metabarcoding. (c) Relative abundance (% reads) of the plant species detected in the third type of mock controls (i.e., genomic DNA pooled from different species of two genus) through *rbcL* and *ITS2* metabarcoding. Detailed list of plant species used in each mock control described in Fig. 2. Abbreviations are mentioned in Fig. 3.

**Table S1** PCR amplification of 45 medicinal plant species with new *rbcL* and *ITS2* metabarcode.

| **Plant species** | ***rbcL*** | ***ITS2*** |
| --- | --- | --- |
| *Aegle marmelos* | + | + |
| *Ailanthus excelsa* | + | - |
| *Aloe vera* | + | + |
| *Andrographis paniculata* | + | - |
| *Asparagua dumosus* | + | + |
| *Asparagus adscendens* | + | + |
| *Asparagus racemosus* | + | + |
| *Azadirachta indica* | + | + |
| *Bacopa monnieri* | + | + |
| *Carica papaya* | + | + |
| *Cassia angustifolia* | + | + |
| *Cassia tora* | + | + |
| *Centella asiatica* | + | + |
| *Eclipta alba* | + | + |
| *Feronia limonia* | + | + |
| *Gossypium herbaceum* | + | + |
| *Hygrophila auriculata* | + | + |
| *Ipomoea batatas* | + | + |
| *Justicia adhatoda syn Adhatoda vasica* | + | - |
| *Lagerstroemia speciosa* | + | + |
| *Lantana camara* | + | + |
| *Melia azedarach* | + | + |
| *Mentha arvensis* | + | + |
| *Ocimum basilicum* | + | + |
| *Ocimum canum* | + | - |
| *Ocimum gratissimum* | + | + |
| *Ocimum tenuiflorum syn. O. sanctum* | + | - |
| *Phyllanthus amarus* | + | + |
| *Phyllanthus emblica* | + | + |
| *Phyllanthus madresapatensis* | + | + |
| *Piper betle* | + | + |
| *Piper longum* | + | + |
| *Pipper arabarum* | + | + |
| *Pipper nigrum* | + | + |
| *Plantago ovata* | + | + |
| *Senna alexandrina* | + | + |
| *Senna occidentalis* | + | + |
| *Solanum nigrum* | + | + |
| *Terminalia arjuna* | + | + |
| *Terminalia bellerica* | + | + |
| *Terminalia chebula* | + | + |
| *Tribulus terrestris* | + | + |
| *Vitex negundo* | + | + |
| *Withania somnifera* | + | + |
| *Zingiber officinale* | + | + |

**Table S2** Metadata of mock controls.

| **Metabarcode** | **Control of mock description** | **Sample ID** | **Reads before filtering** | **Cumulative reads per control** | **Reads after filtering** | **Percentage of analysed reads from filtered reads^#^** |
| --- | --- | --- | --- | --- | --- | --- |
| ***rbcL*** | Control 1: Genomic DNA pool of different genus | 5P | 2877 | 18657 | 2417 | 84.0 |
|  |  | 10P | 7993 |  | 7679 | 96.1 |
|  |  | 15P | 7787 |  | 6742 | 86.6 |
|  | Control 2: Simulated plant biomass controls (blended formulations) | 5S | 14695 | 42140 | 12406 | 84.4 |
|  |  | 10S | 14183 |  | 12221 | 86.2 |
|  |  | 15S | 13262 |  | 11402 | 86.0 |
|  | Control 3: Genomic DNA pooled from different species of the genus | 6SP | 8754 | 10265 | 7993 | 91.3 |
|  |  | 7SP | 1511 |  | 1336 | 88.4 |
| ***ITS2*** | Control 1: Genomic DNA pool of different genus | 5P | 113942 | 452380 | 90900 | 79.8 |
|  |  | 10P | 140647 |  | 83209 | 59.2 |
|  |  | 15P | 197791 |  | 134011 | 67.8 |
|  | Control 2: Simulated plant biomass controls (blended formulations) | 5S | 171919 | 334017 | 94671 | 55.1 |
|  |  | 10S | 88165 |  | 40403 | 45.8 |
|  |  | 15S | 73933 |  | 33001 | 44.6 |
|  | Control 3: Genomic DNA pooled from different species of the genus | 6SP | 63249 | 164358 | 38013 | 60.1 |
|  |  | 7SP | 101109 |  | 60328 | 59.7 |

**Table S3** Metadata of single drugs.

| **Metabarcode** | **Single drug description** | **Sample ID** | **Total Reads obtained** | **cumulative total reads** | **Reads after filtering** | **Percentage of analysed reads from filtered reads** |
| --- | --- | --- | --- | --- | --- | --- |
| ***rbcL*** | Tulsi churna | 1 | 5645 | 128540 | 5362 | 89.2 |
|  |  | 2 | 8534 |  | 8162 | 80.5 |
|  |  | 3 | 11571 |  | 8537 | 69.1 |
|  |  | 4 | 7984 |  | 7563 | 85.2 |
|  | Gokharu churna | 5 | 7993 |  | 6208 | 78.8 |
|  |  | 6 | 9189 |  | 6468 | 78.0 |
|  |  | 7 | 3871 |  | 3238 | 83.5 |
|  |  | 8 | 3508 |  | 2746 | 74.6 |
|  |  | 9 | 5389 |  | 4264 | 79.9 |
|  | shatavari churna | 10 | 6225 |  | 5371 | 91.1 |
|  |  | 11 | 19350 |  | 17666 | 96.0 |
|  |  | 12 | 21915 |  | 19996 | 95.8 |
|  | Vasa churna | 13 | 4080 |  | 3549 | 74.0 |
|  |  | 14 | 2838 |  | 2535 | 87.6 |
|  | Bhurgra churna | 15 | 4471 |  | 3060 | 78.3 |
|  | Ashwagandha churna | 16 | 4042 |  | 3570 | 86.6 |
|  | Arjuna churna | 17 | 1935 |  | 982 | 75.4 |
| ***ITS2*** | Tulsi churna | 1 | 9034 | 1211935 | 4913 | 28.3 |
|  |  | 2 | 19193 |  | 17427 | 73.6 |
|  |  | 3 | 30858 |  | 21491 | 57.8 |
|  |  | 4 | 43781 |  | 33716 | 68.7 |
|  | Gokharu churna | 5 | 60107 |  | 42606 | 89.0 |
|  |  | 6 | 67986 |  | 31295 | 87.9 |
|  |  | 7 | 14625 |  | 3736 | 51.4 |
|  |  | 8 | 58734 |  | 46804 | 84.6 |
|  |  | 9 | 89731 |  | 30965 | 88.9 |
|  | Shatavari churna | 10 | 102215 |  | 23772 | 49.9 |
|  |  | 11 | 167121 |  | 109646 | 91.4 |
|  |  | 12 | 70460 |  | 44038 | 95.5 |
|  | Vasa churna | 13 | 110041 |  | 20587 | 58.8 |
|  |  | 14 | 62479 |  | 48048 | 74.6 |
|  | Bhurgra churna | 15 | 177487 |  | 135568 | 90.7 |
|  | Ashwagandha churna | 16 | 102700 |  | 14221 | 75.9 |
|  | Arjuna churna | 17 | 25383 |  | 21114 | 82.7 |

**Table S4** Relative abundance (% reads) of non-targeted plant species detected within single drugs.

| **Metabarcode** | **Single drug description** | **Sample ID** | **Non targeted reads** | **Percentage of reads from total analysed reads^#^** |
| --- | --- | --- | --- | --- |
| ***rbcL*** | Tulsi churna | 1 | *Others* | 0.2 |
|  |  | 2 | *Biancaea sappan* | 14.4 |
|  |  |  | *Rotala rotundifolia* | 1.6 |
|  |  |  | *Azadirachta indica* | 1.3 |
|  |  |  | *Others* | 0.6 |
|  | Gokharu churna | 5 | *Biancaea sappan* | 2.9 |
|  |  |  | *Apium graveolens* | 3.0 |
|  |  |  | *Others* | 1.3 |
|  |  | 6 | *Ligusticum jeholense* | 1.3 |
|  |  |  | *Buddleja alternifolia* | 94.5 |
|  |  |  | *Others* | 4.2 |
|  |  | 7 | *Abutilon theophrasti* | 1.2 |
|  |  |  | *Others* | 1.4 |
|  |  | 8 | *Biancaea sappan* | 12.7 |
|  |  |  | *Canavalia cathartica* | 24.0 |
|  |  |  | *Others* | 2.9 |
|  |  | 9 | *Others* | 1.7 |
|  | Shatavari churna | 10 | *Others* | 0.3 |
|  |  | 11 | *Others* | 0.4 |
|  | Bhurgraj churna | 15 | *Others* | 1.2 |
|  | Ashwagandha churna | 16 | *Physochlaina physaloides* | 17.2 |
|  |  |  | *Abutilon theophrasti* | 2.5 |
|  |  |  | *Others* | 4.9 |
| ***ITS2*** | Tulsi churna | 1 | *Uncultured eukaryote* | 41.9 |
|  |  |  | *Olea europaea* | 23.7 |
|  |  |  | *Lawsonia inermis* | 5.0 |
|  |  |  | *Lamiaceae sp.* | 2.9 |
|  |  |  | *Indigofera tinctoria* | 2.5 |
|  |  |  | *Ocimum africanum* | 2.4 |
|  |  |  | *Alternanthera halimifolia* | 2.4 |
|  |  |  | *Lobelia cliffortiana* | 2.1 |
|  |  |  | *Trifolium alexandrinum* | 1.6 |
|  |  |  | *Pentanema indicum* | 1.3 |
|  |  |  | *Others* | 2.5 |
|  |  | 2 | *Senna alexandrina* | 95.8 |
|  |  |  | *Senna italica* | 2.5 |
|  |  |  | *Lawsonia inermis* | 1.2 |
|  |  |  | *Others* | 0.5 |
|  |  | 3 | *Glycyrrhiza glabra* | 26.8 |
|  |  |  | *Cuminum cyminum* | 14.7 |
|  |  |  | *Lobelia cliffortiana* | 6.4 |
|  |  |  | *Foeniculum vulgare* | 4.3 |
|  |  |  | *Euphorbia heterophylla* | 4.1 |
|  |  |  | *Amaranthus viridis* | 4.1 |
|  |  |  | *Chloris barbata* | 3.5 |
|  |  |  | *Acalypha indica* | 2.5 |
|  |  |  | *Abutilon pannosum* | 2.3 |
|  |  |  | *Corynandra felina* | 2.1 |
|  |  |  | *Parthenium hysterophorus* | 2.0 |
|  |  |  | *Croton curiosus* | 2.0 |
|  |  |  | *Artemisia salsoloides* | 1.6 |
|  |  |  | *Eclipta prostrata* | 1.5 |
|  |  |  | *Alysicarpus scariosus* | 1.4 |
|  |  |  | *Boerhavia erecta* | 1.3 |
|  |  |  | *Indigofera linnaei* | 1.2 |
|  |  |  | *Panicum grumosum* | 1.1 |
|  |  |  | *Others* | 17.3 |
|  |  | 4 | *Parthenium hysterophorus* | 49.8 |
|  |  |  | *Cuscuta campestris* | 21.2 |
|  |  |  | *Dactyloctenium aegyptium* | 5.3 |
|  |  |  | *Sida acuta* | 3.4 |
|  |  |  | *Cuscuta pentagona* | 2.5 |
|  |  |  | *Amaranthus viridis* | 1.4 |
|  |  |  | *Eclipta prostrata* | 1.4 |
|  |  |  | *Setaria verticillata* | 1.2 |
|  |  |  | *Abutilon indicum* | 1.2 |
|  |  |  | *Canavalia vitiensis* | 1.0 |
|  |  |  | *Others* | 11.5 |
|  | Gokharu churna | 5 | *Others* | 1.0 |
|  |  | 6 | *Senna alexandrina* | 1.5 |
|  |  |  | *Glycyrrhiza glabra* | 1.9 |
|  |  |  | *Cuminum cyminum* | 4.3 |
|  |  |  | *Foeniculum vulgare* | 5.3 |
|  |  |  | *Moringa oleifera* | 6.0 |
|  |  |  | *Others* | 4.6 |
|  |  | 7 | *Others* | -0.6 |
|  |  | 8 | *Canavalia dictyota* | 1.7 |
|  |  |  | *Canavalia gladiata* | 3.9 |
|  |  |  | *Senna alexandrina* | 4.3 |
|  |  |  | *Others* | 1.7 |
|  | Shatavari churna | 9 | *Cuminum cyminum* | 10.0 |
|  |  |  | *Carum carvi* | 5.9 |
|  |  |  | *Tribulus pentandrus* | 3.5 |
|  |  |  | *Trachyspermum ammi* | 3.2 |
|  |  |  | *Wallemia sp.* | 2.9 |
|  |  |  | *Uncultured fungus* | 2.7 |
|  |  |  | *Uncultured eukaryote* | 1.4 |
|  |  |  | *Others* | 2.7 |
|  |  | 10 | *Cuminum cyminum* | 25.1 |
|  |  |  | *Foeniculum vulgare* | 3.9 |
|  |  |  | *Senna alexandrina* | 2.2 |
|  |  |  | *Others* | 4.2 |
|  |  | 11 | *Tribulus pentandrus* | 24.0 |
|  |  |  | *Baccharoides anthelmintica* | 3.7 |
|  |  |  | *Hibiscus micranthus* | 2.1 |
|  |  |  | *Others* | 8.5 |
|  | Bhurgraj churna | 15 | *Others* | 4.3 |
|  | Ashwgandha churna | 16 | *Abutilon indicum* | 44.5 |
|  |  |  | *Others* | 8.2 |

^#^Plant species that comprised <1% reads are represented as others

**Table S5** Metadata of polyherbal formulations.

| **Metabarcode** | **Single drug description** | **Sample ID** | **Reads before filtering** | **cumulative reads** | **Reads after filtering** | **Percentage of analysed reads from filtered reads** |
| --- | --- | --- | --- | --- | --- | --- |
| ***rbcL*** | Trikatu churna | 18 | 2178 | 53087 | 1852 | 71.9 |
|  |  | 19 | 1080 |  | 624 | 50.3 |
|  |  | 20 | 3078 |  | 2268 | 72.4 |
|  | Sitopaladi churna | 21 | 2564 |  | 1857 | 73.1 |
|  |  | 22 | 5930 |  | 4336 | 76.7 |
|  |  | 23 | 6393 |  | 4165 | 71.8 |
|  | Rasayana churna | 24 | 3148 |  | 2070 | 90.9 |
|  |  | 25 | 6049 |  | 4556 | 78.9 |
|  |  | 26 | 4110 |  | 2916 | 71.7 |
|  |  | 27 | 3117 |  | 2621 | 51.2 |
|  | Hingwashtak churna | 28 | 4052 |  | 2293 | 81.7 |
|  |  | 29 | 3513 |  | 2873 | 79.5 |
|  |  | 30 | 4166 |  | 3536 | 79.5 |
|  |  | 31 | 2630 |  | 2266 | 80.7 |
|  | Talisadi churna | 32 | 1079 |  | 847 | 69.9 |
| ***ITS2*** | Trikatu churna | 18 | 151194 | 1429238 | 106886 | 84.4 |
|  |  | 19 | 14661 |  | 8610 | 59.5 |
|  |  | 20 | 11774 |  | 5497 | 52.2 |
|  | Sitopaladi churna | 21 | 31624 |  | 24503 | 85.8 |
|  |  | 22 | 38070 |  | 26608 | 75.5 |
|  |  | 23 | 43870 |  | 26608 | 66.2 |
|  | Rasayana churna | 24 | 198979 |  | 162279 | 94.8 |
|  |  | 25 | 52189 |  | 40712 | 87.4 |
|  |  | 26 | 65524 |  | 52564 | 89.7 |
|  |  | 27 | 32193 |  | 25781 | 85.0 |
|  | Hingwashtak churna | 28 | 37952 |  | 34096 | 81.1 |
|  |  | 29 | 201382 |  | 128303 | 88.4 |
|  |  | 30 | 261442 |  | 216913 | 98.3 |
|  |  | 31 | 225768 |  | 108909 | 98.6 |
|  | Talisadi churna | 32 | 62616 |  | 49231 | 86.4 |

**Table S6** Percentage reads of non-targeted plant species detected within polyherbal formulations.

| **Metabarcode** | **Single drug description** | **Sample ID** | **Non targeted reads** | **Percentage of reads from total analysed reads**^#^ |
| --- | --- | --- | --- | --- |
| ***rbcL*** | Trikatu churna | 19 | *Trigonella foenum-graecum* | 2.9 |
|  |  | 20 | *Halopegia azurea* | 1.0 |
|  |  |  | *Others* | 0.8 |
|  | Sitopaladi churna | 21 | *Others* | 1.8 |
|  |  | 22 | *Others* | 3.2 |
|  |  | 23 | *Others* | 0.8 |
|  | Rasayana churna | 25 | *Others* | 0.2 |
|  |  | 27 | *Others* | 1.4 |
|  | Hingvashtak churna | 28 | *Others* | 0.6 |
|  |  | 29 | *Others* | 1.3 |
|  |  | 30 | *Others* | 4.0 |
|  |  | 31 | *Melilotus albus* | 2.0 |
|  |  |  | *Caucalis platycarpos* | 1.9 |
|  |  |  | *Others* | 0.4 |
|  | Talisadya churna | 32 | *Others* | 0.8 |
| ***ITS2*** | Trikatu churna | 18 | *Others* | 0.6 |
|  |  | 19 | *Indigofera cordifolia* | 12.8 |
|  |  |  | *Trigonella foenumgraecum* | 11.0 |
|  |  |  | *Uncultured eukaryote* | 5.3 |
|  |  |  | *Foeniculum vulgare* | 4.8 |
|  |  |  | *Cuminum cyminum* | 1.9 |
|  |  |  | *Ocimum tenuiflorum* | 1.1 |
|  |  |  | *Others* | 3.2 |
|  |  | 20 | *Ocimum tenuiflorum* | 15.3 |
|  |  |  | *Uncultured eukaryote* | 7.9 |
|  |  |  | *Others* | 1.2 |
|  | Sitopaladi churna | 21 | *Foeniculum vulgare* | 20.4 |
|  |  |  | *Tribulus pentandrus* | 2.9 |
|  |  |  | *Cuminum cyminum* | 1.8 |
|  |  |  | *Others* | 2.5 |
|  |  | 22 | *Foeniculum vulgare* | 45.9 |
|  |  | 22 | *Mucuna pruriens* | 1.3 |
|  |  |  | *Senna alexandrina* | 1.1 |
|  |  |  | *Others* | 1.4 |
|  |  | 23 | *Cuminum cyminum* | 1.6 |
|  |  |  | *Others* | 1.4 |
|  | Rasayana churna | 24 | *Others* | 0.1 |
|  |  | 26 | *Others* | 0.4 |
|  |  | 27 | *Foeniculum vulgare* | 2.0 |
|  |  |  | *Others* | 0.3 |
|  | Hingvashtak churna | 28 | *Others* | 0.4 |
|  |  | 29 | *Foeniculum vulgare* | 3.3 |
|  |  |  | *Others* | 1.9 |
|  |  | 30 | *Seseli diffusum* | 4.5 |
|  |  | 31 | *Others* | 0.4 |
|  | Talisadya churna | 32 | *Olea europaea* | 11.8 |
|  |  |  | *Others* | 1.1 |

^#^Plant species that comprised <1% reads are included in others
